# Tumor-wide RNA splicing aberrations generate immunogenic public neoantigens

**DOI:** 10.1101/2023.10.19.563178

**Authors:** Darwin W. Kwok, Nicholas O. Stevers, Takahide Nejo, Lee H. Chen, Inaki Etxeberria, Jangham Jung, Kaori Okada, Maggie Colton Cove, Senthilnath Lakshmanachetty, Marco Gallus, Abhilash Barpanda, Chibo Hong, Gary K.L. Chan, Samuel H. Wu, Emilio Ramos, Akane Yamamichi, Jerry Liu, Payal Watchmaker, Hirokazu Ogino, Atsuro Saijo, Aidan Du, Nadia Grishanina, James Woo, Aaron Diaz, Susan M. Chang, Joanna J. Phillips, Arun P. Wiita, Christopher A. Klebanoff, Joseph F. Costello, Hideho Okada

## Abstract

T-cell-mediated immunotherapies are limited by the extent to which cancer-specific antigens are homogenously expressed throughout a tumor. We reasoned that recurrent splicing aberrations in cancer represent a potential source of tumor-wide and public neoantigens, and to test this possibility, we developed a novel pipeline for identifying neojunctions expressed uniformly within a tumor across diverse cancer types. Our analyses revealed multiple neojunctions that recur across patients and either exhibited intratumor heterogeneity or, in some cases, were tumor-wide. We identified CD8+ T-cell clones specific for neoantigens derived from tumor-wide and conserved neojunctions in *GNAS* and *RPL22*, respectively. TCR-engineered CD8^+^ T-cells targeting these mutations conferred neoantigen-specific tumor cell eradication. Furthermore, we revealed that cancer-specific dysregulation in splicing factor expression leads to recurrent neojunction expression. Together, these data reveal that a subset of neojunctions are both intratumorally conserved and public, providing the molecular basis for novel T-cell-based immunotherapies that address intratumoral heterogeneity.

## INTRODUCTION

Immunotherapy is a promising strategy for cancer treatment, showing remarkable survival benefit in various disease types.^1–6^ However, many tumors evade eradication due to intratumoral heterogeneity (ITH).^7–12^ ITH refers to the diversification of a tumor’s cellular and genetic landscape, which enables subsets of tumor cells to resist or evade eradication.^13–15^ Despite the success of immunotherapy in tumors with high immune infiltration and mutational load,^16–18^ cancer types with extensive ITH or lower mutational burdens are more resistant.^19–22^ Therefore, it is critical to expand the targetable antigen repertoire for T-cell-based therapies and comprehensively consider ITH in developing novel immunotherapy strategies for improving outcomes.

Current immunotherapies targeting tumor-specific antigens (TSAs) have focused on peptides that arise from somatic mutations in canonical coding regions of the genome.^6,23^ Due to its reliance on DNA-based nonsynonymous mutations, this approach yields limited targets in tumors with low mutational burdens.^24,25^ To expand the repertoire of potential immunotherapeutic antigens, recent studies have explored aberrant splicing events across multiple cancers as an additional source of TSAs.^26,27^ These cancer-specific splicing events, otherwise known as neojunctions, are prevalent in cancer cells and capable of generating novel TSAs that potentiate CD8^+^ T-cell-mediated expansion and responses in select cancer types.^26,28,30,34–38^ Nevertheless, to date, no study has examined the conservation of neojunctions across whole tumors, and therefore, those identified neojunction-derived targets face similar therapeutic barriers due to ITH.

Herein, we sought to fill this gap in knowledge by investigating the clonality of neojunctions across cancer types to identify public, tumor-wide neojunction-derived TSAs. To accomplish this goal, we generated a high-throughput and comprehensive spatial splicing-derived neoantigen identifier pipeline for characterizing neojunctions found in multiple intratumoral sites (spatially conserved) by systematically mapping RNA splicing junctions across distinct regions within the same tumor **(Figure 1).** This pipeline identified immunogenic neojunction-derived TSAs, and further revealed that public tumor-wide neojunctions generate proteolytically-processed and major histocompatibility class-I (MHC-I)-bound TSAs. Recognition of these antigens by TSA-specific CD8^+^ T-cells induced T-cell receptor (TCR) signaling and tumor cell tumor killing. Together, these findings highlight the potential of targeting public neojunction-derived TSAs as a novel class of “off-the-shelf” cancer immunotherapies offering a promising avenue for improving cancer treatment.

**Figure 1:**
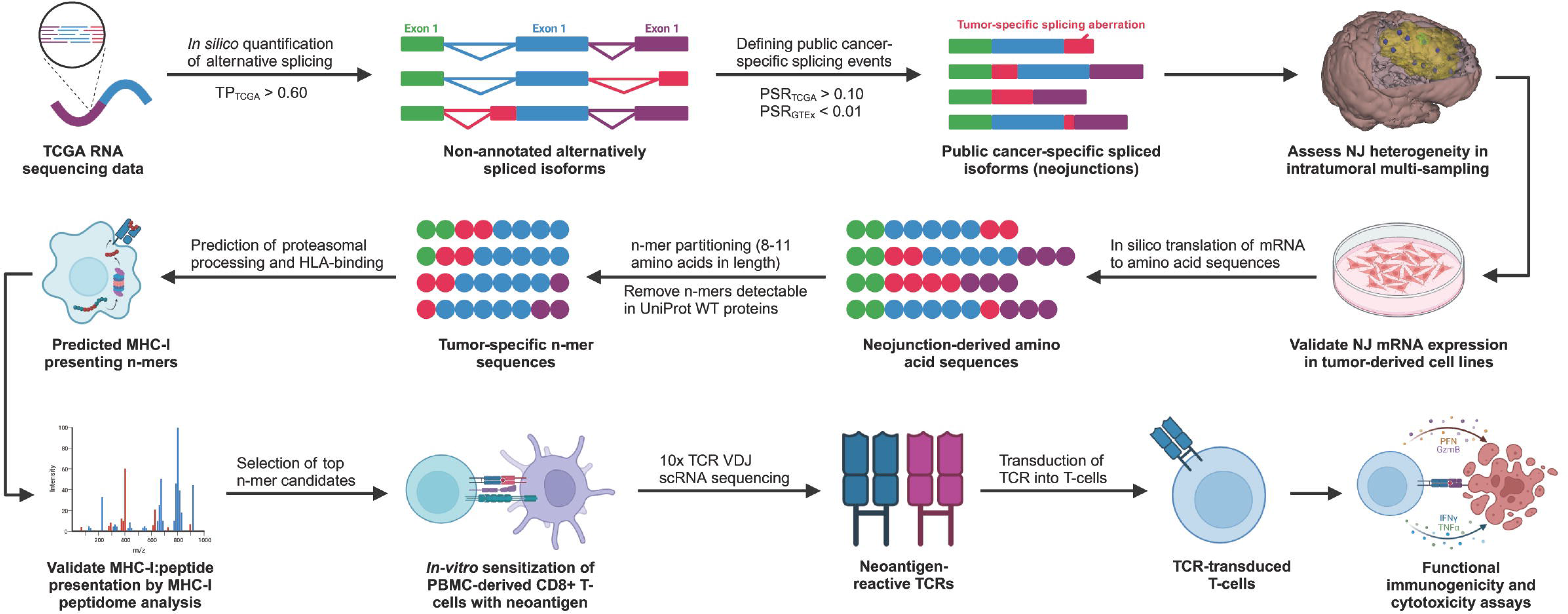
Novel neoantigen discovery pipeline for discovering public and tumor-wide targets for cellular immunotherapy across various cancers. TCGA RNA sequencing data across multiple cancers (*n*=11) were analyzed for non-annotated, protein-coding, and cancer-specific splicing junctions (GTEx positive sample rate < 1%; neojunctions). Interpatiently conserved (TCGA positive sample rate ≥ 10%; public neojunctions) were retained for downstream analysis of intratumoral heterogeneity (ITH). Tumors with sequencing data extracted from multiple intratumoral regions were used to evaluate each public neojunction’s ITH. Independent prediction algorithms were used to assess proteasomal processing and MHC-I binding of peptide sequences translated from public, intratumorally conserved neojunctions. The expression of these neojunctions and their peptide derivatives were validated by RNA sequencing and mass spectrometry analysis of patient-derived tumor samples and cell lines. T-cell receptors (TCRs) were cloned and characterized for top predicted candidates through *in vitro* sensitization of PBMC-derived CD8^+^ T-cells against the corresponding neoantigen-pulsed antigen presenting cells and subsequent 10x V(D)J single-cell sequencing. Transduction of these neoantigen-reactive TCR sequences in TCR-null Jurkat76/CD8 cells and PBMC-derived CD8^+^ T-cells allowed the demonstration of neoantigen-specific immunogenicity and tumor-specific killing.

## RESULTS

### Characterization of public, pan-cancer neojunctions

To investigate the expression of neojunctions, we first utilized RNA sequencing (RNA-seq) data from The Cancer Genome Atlas (TCGA) to identify nonannotated junction reads across various cancer types **(Figure 2A).** Our analysis was also applied to tumors with RNA-seq from multiple spatially mapped tumor samples **(Figures 2B, S1)**. These included sequencing data sets from glioblastoma, low-grade glioma, lung adenocarcinoma, lung squamous cell carcinoma, mesothelioma, liver hepatocellular carcinoma, stomach adenocarcinoma, kidney renal clear cell carcinoma, kidney renal papillary cell carcinoma, chromophobe renal cell carcinoma, colon adenocarcinoma, and prostate adenocarcinoma. To minimize the confounding issue of non-malignant cell admixture, only samples with inferred tumor purities of 60% or higher were included^29,31^ **(Figure 2C)**, and protein-coding, nonannotated junctions were identified **(Figure S2A).** A junction’s positive sample rate (PSR) represents the percentage of samples within a cohort expressing the neojunction with a junction read frequency ≥ 1% compared to the canonical splicing junction. By focusing on robustly expressed neojunctions that are shared among patients, we filtered for public neojunctions that demonstrate elevated PSRs within each tumor type (PSR_TCGA_ ≥ 10%) **(Figure 2D, S2B)**. Following previous neojunction nomenclature,^26^ we selected for splicing events expressed with a PSR lower than 1% in normal tissue RNA-seq data from the Genotype-Tissue Expression (GTEx) project (n=9166; PSR_GTEx_ < 1%) **(Figures S2C)**. We identified an average of 373 neojunctions per disease across TCGA tumor types, of which 202 were public **(Figure 2E, Supplementary Table 1)** and expressed at similar frequencies **(Figure 2F)**. Further characterization of public neojunctions across tumor types demonstrated a predominance of alternative 3’ and 5’ splice site changes **(Figure 2G)**, and the proportion of frame shift-generating neojunctions was consistent across tumors **(Figure 2H)**. The large majority of neojunctions in our pan-cancer analysis are expressed across multiple disease types. Unbiased hierarchical clustering revealed that cases from the same tumor type relatively clustered together, which indicates that the neojunction expression profile remained consistent across patients of the same disease type. However, subsets of neojunctions were shown by this same study to be expressed across multiple tumor types, indicating the presence of potential targets that span various diseases **(Figure S2D)**. Focusing on a small subset of these neojunctions **(Figure 2I),** we identified neojunctions that were expressed in most, if not all, tumor subtypes. These results indicate a significant conservation of neojunction expression among patients across multiple cancer types, suggesting the possibility of a pan-cancer immunotherapy targeting products of aberrant splicing.

**Figure 2:**
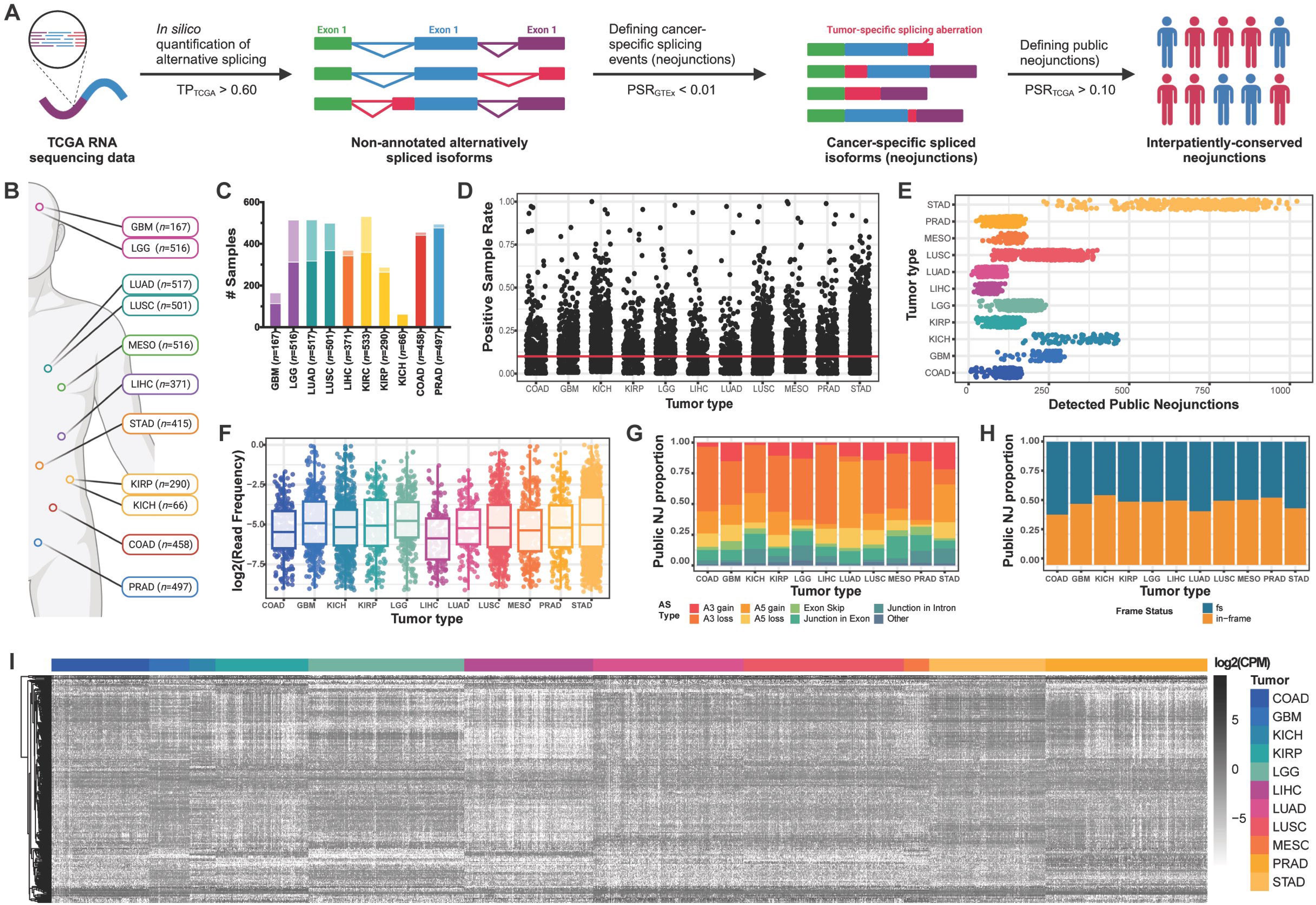
Public putative neojunctions are characterized across multiple cancer types. **A.** Pipeline for identifying putative, spatially conserved, cancer-specific alternative splicing events (neojunctions) from RNA-sequencing data collected from The Cancer Genome Atlas (TCGA). **B.** TCGA tumor sets with corresponding multi-biopsy RNA sequencing data available for analysis. Disease types were selected based on the availability of publicly available data sets that included multi-site sampling within the same tumor, which included glioblastoma (GBM; n=167), low-grade glioma (LGG; n=516), lung adenocarcinoma (LUAD, n=517), lung squamous cell carcinoma (LUSC, n=501), mesothelioma (MESO, n=516), liver hepatocellular carcinoma (LIHC, n=371), stomach adenocarcinoma (STAD, n=415), kidney renal clear cell carcinoma (KIRC; n=533), kidney renal papillary cell carcinoma (KIRP; n=290), kidney chromophobe (KICH, n=66), colon adenocarcinoma (COAD; n=458), and prostate adenocarcinoma (PRAD; n=497). **C**. Tumor purity of TCGA tumor samples. Samples with confirmed tumor purity of 60% and above (solid fraction) were retained for downstream analysis. **D.** Interpatient frequency (positive sample rate; PSR) of putative neojunctions identified in each tumor type. Public neojunctions are defined as those with a PSR ≥ 10% (red line). **E.** Total public neojunctions detected per sample across tumor types. **F.** log_2_(read frequency) of public neojunctions across tumor types. **G-H.** Distribution of public neojunctions based on splice types (**G**) and frame-shift status (**H**). **I.** Expression of all public neojunctions (log_2_(CPM)) across all studied TCGA tumor types.

### Neojunctions exhibit intratumor heterogeneity

Immunotherapy approaches targeting a single TSA might be inadequate to eradicate tumors because of the outgrowth of the non-antigenic population within the host. In the case of targeted T-cell therapies, this underscores the importance of focusing on multiple neoantigens shared by the entire tumor to prevent antigenic evolution.^32,33^ Neojunctions can generate immunogenic antigens,^34–36,38^ and to that end, we explored the ITH of public neojunctions by filtering for neojunction reads across multiple samples from the same tumor **(Figure 3A)**. RNA-seq data derived from multiple intratumor samples in prostate,^39^ liver,^40–43^ colon,^40,44^ stomach,^40^ kidney,^40^ and lung^45,46^ were analyzed to investigate whether public neojunctions are present across the tumor landscape in various cancer types **(Figures S1)**. This analysis revealed public neojunctions expressed in multiple samples, a subset of which were also found tumor-wide across all biopsies **(Figure 3B, S3A-S3C).** The identification of public neojunctions expressed tumor-wide suggests a potential immunotherapy approach that is capable of targeting every neoplastic compartment. In datasets containing multiple cases with multi-site sampling, public spatially conserved neojunctions were expressed readily across all cases, further indicating ideal neojunction targets that are expressed both intratumorally and interpatiently. **(Figure 3C)**.

**Figure 3:**
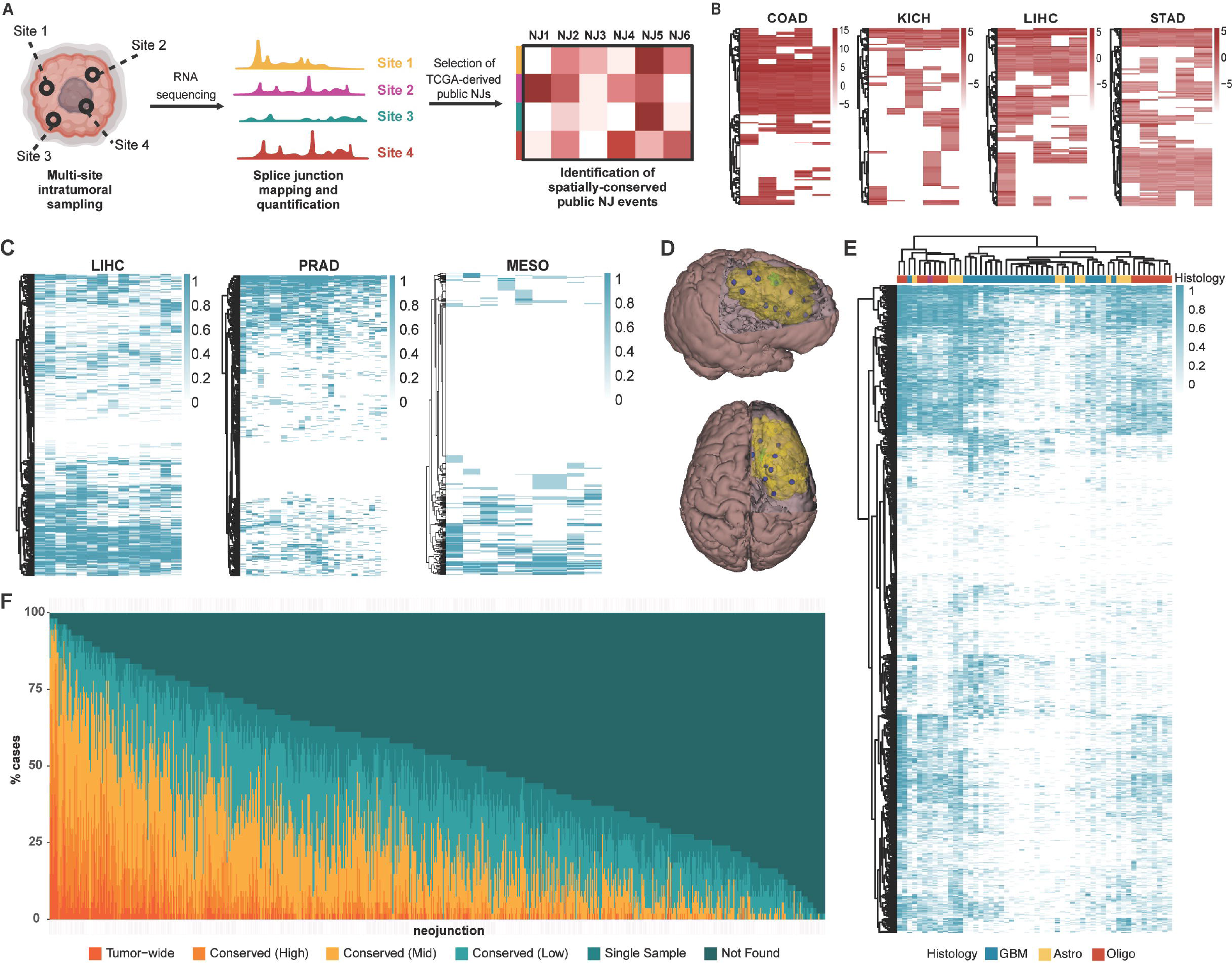
Subsets of neojunctions are expressed tumor-wide. **A.** Overview of the tumor-wide characterization of neojunctions by investigating RNA sequencing of multiple intratumoral regions in various cancers. **B.** Heatmaps representing log_2_(CPM) of neojunctions (rows) across five samples within the same tumor (columns) in colon cancer, kidney cancer, liver cancer, and stomach cancer. Neojunctions found across all five intratumoral samples are annotated in yellow. **C.** Heatmap illustrating the number of biopsies within the same tumor sample (columns) that has detectable expression of neojunctions (rows) in liver cancer (left), prostate cancer (center), and mesothelioma (right). The intensity of each cell indicates the proportion of regions within the same tumor that has putative expression of each neojunction, with intensity of 1 representing a neojunction expressed tumor-wide within a corresponding patient. **D.** 3-D models of the brain and tumor (yellow) derived from patient 470 (P470). Approximately 10 spatially mapped and maximally distanced biopsies (blue) were taken within each tumor (refer to Supplementary Video 1). Whole-exome sequencing, RNA sequencing, and further analyses were conducted on each of these regions. **E.** Heatmap illustrating the number of biopsies within the same tumor sample (columns) that has detectable expression of neojunctions (rows) in across various glioma subtypes, *IDH1*wt (blue), *IDH1*mut-A (yellow), and *IDH1*mut-O (red). The intensity of each cell indicates the percentage of regions within the same tumor that has putative expression of each neojunction. **F.** Distribution of glioma-specific neojunctions (*n*=789, columns) based on the their intratumoral heterogeneity across patients. Neojunctions are considered tumor-wide if they are found across 100% of intratumor samples within the same patient, highly conserved if they are found in > 70% of samples but not 100% of samples, moderately conserved if they are found in > 30% of samples but ≤ 70% of samples, or lowly conserved if found in at least two samples but ≤ 30% of samples.

Of the cancer types analyzed in our study, gliomas have a notoriously greater degree of ITH that further complicates immunotherapeutic applications. The heterogenous nature of gliomas is evidenced by studies using single-cell RNA-seq data from isocitrate dehydrogenase 1 (*IDH1*)-wild type (*IDH1*wt) and *IDH1*-mutant (*IDH1*mut) gliomas which revealed numerous subclones with branched evolutionary trajectories.^7,8,47,48^ While multi-site RNA-seq data from a handful of intratumoral biopsies are publicly available, it is difficult to conclude whether a few samples are sufficient in fully recapitulating a tumor’s transcriptomic landscape, especially in highly heterogeneous tumors, such as gliomas.^7,49^ As such, we have utilized our own dataset to significantly expand the number of intratumoral biopsies analyzed across the three main glioma subtypes.^50^ Building upon the previous dataset, we evaluated approximately 10 maximally distanced, spatially-mapped biopsies from a total of 56 tumors, including all three glioma subtypes, for whole-exome and RNA-seq **(Figure 3D)**. Similar to multi-region studies in other tumor types, we detected expression of neojunctions intratumorally across multiple patients. Hierarchal clustering revealed neojunction subsets tend to associate with either *IDH1*mut subtypes or *IDH1*wt subtypes **(Figure 3E)**, and interestingly, we noticed that *IDH1*mut gliomas had significantly greater number of neojunctions that were expressed tumor-wide when compared to *IDH1*wt gliomas. When investigating putative expression of the 789 total neojunctions initially identified across both LGG and GBM TCGA cases, 774 (98.10%) and 547 (69.33%) neojunctions were detectable in more than one region within at least one tumor and 10% of a glioma subtype cohort, respectively **(Figure 3F)**. This indicates that the majority of the public neojunctions identified from the TCGA LGG/GBM analysis are consistently expressed in multiple tumor regions across our comprehensive patient database. This finding is significant in demonstrating that targeting these neojunctions in combination can offer a therapeutic avenue that covers the entire tumor landscape. Tumor-wide neojunctions detectable in all biopsies within a single tumor exhibited the pinnacle characteristics of intratumor conservation. Remarkably, 280 (34.49%) of the neojunctions were robustly expressed tumor-wide within at least one case, Underscoring the feasibility of personalized tumor-wide neojunction-derived immunotherapies. Furthermore, half of these neojunctions were found tumor-wide in two or more cases **(Figure 3G)**, underpinning promising public immunotherapy options that targets tumor-wide neoantigens. Thus, our findings highlight a novel class of neojunctions that appear intratumorally in multiple patients, establishing a promising new repertoire of potential immunologic targets for cancer therapy.

### Public neojunctions are detectable on an RNA and peptide level in patient-derived tumor samples

We next sought to validate whether the expression of public neojunctions and their protein products can be detected in cell line transcriptomic data and patient-derived proteomic data from tumors. Confirming the expression of these neojunctions in cancer cell lines underscores the feasibility of using them as a model for downstream immunogenicity assays. For this analysis, we focused on public neojunctions expressed by gliomas as these tumors display a high degree of ITH, exhibit poor clinical outcomes, and lack potentially curative immunotherapy treatments. We first investigated RNA-seq data of GBM patient-derived xenografts (PDX) (*n*=66)^51^ and LGG (*n*=2) cell lines. RNA-seq data for astrocytoma and oligodendroglioma LGG cell lines were generated in our previous study.^52^ In total, we detected 510 (64.6%) and 767 (97.2%) of our previously characterized public neojunctions expressed at detectable levels in GBM and LGG, respectively **(Figures 4A, 4B).** Bulk RNA-seq has limited reads spanning each splicing aberration. We therefore designed primers spanning a subset of neojunctions and their flanking exons, performed deep amplicon sequencing, and detected the mRNA expression of neojunction-spanning reads expressed within glioma cell lines **(Figure S4H).**

**Figure 4:**
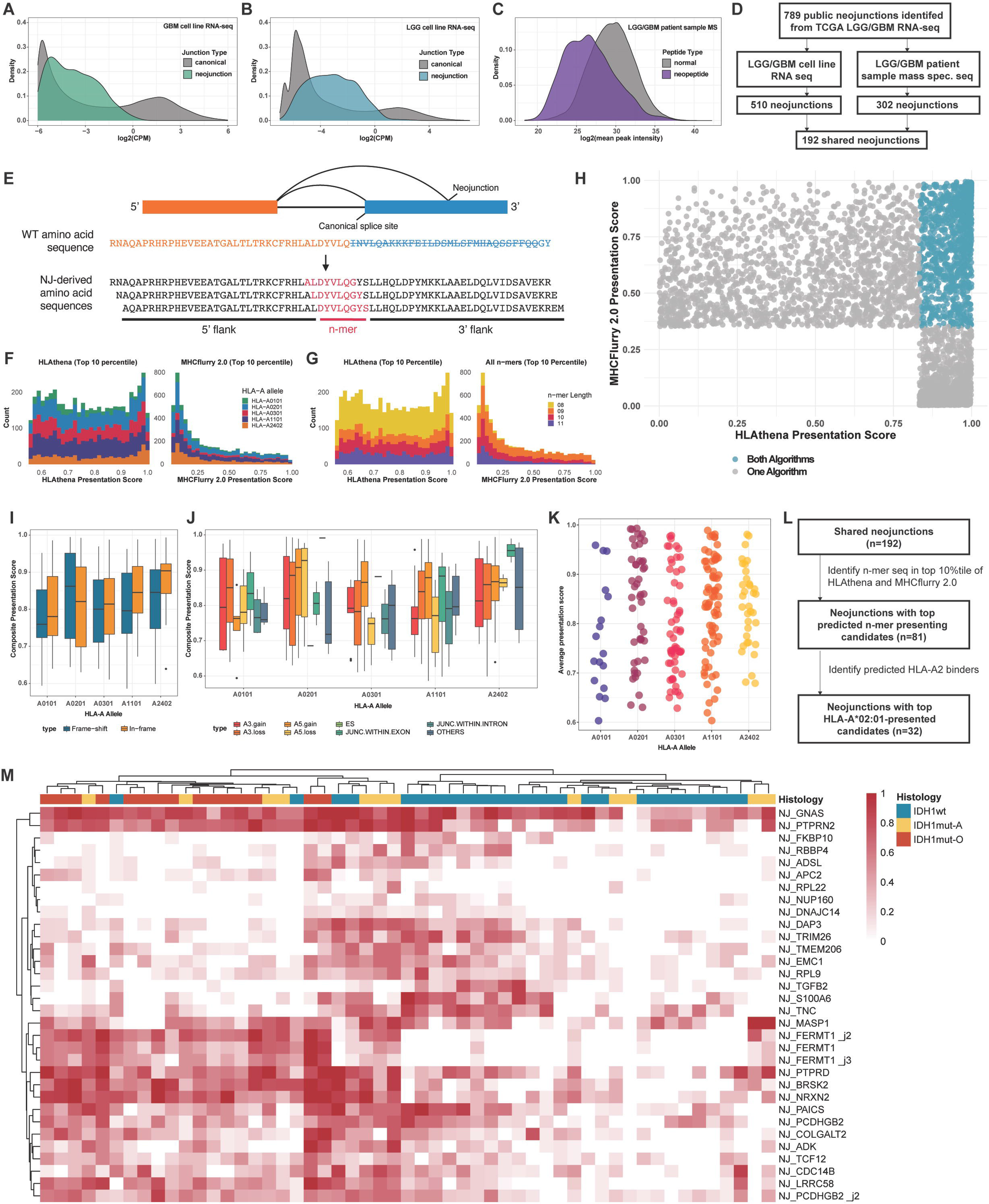
Neojunction-derived neoepitopes are predicted to be processed and presented by MHC-I. **A-B.** Density plots depicting log_2_(CPM) of junction reads derived from RNA-sequencing from patient-derived GBM (**A**) and LGG (**B**) cell lines that validate detectable levels of neojunction expression (colored) compared to the expression of canonical splicing (gray). **C.** Density plot depicting mass spectrometry analysis of multiple publicly-available LGG and GBM data sets (*n*=447). The log2(peak intensity) of detectable neojunction-derived peptides (purple) indicate comparable intensity compared to other endogenous peptides (gray). **D.** Schematic demonstrating the selection of high-confidence neojunctions for downstream analysis. 192 neojunctions were selected as they were found to be expressed in all RNA-seq and MS platforms. **E.** Diagram illustrating the mechanism of neoantigen production by the introduction of a neojunction. Multimer partitioning was employed subsequently to generate a peptide bank for prediction analysis. **F-G.** Histogram illustrating the scores pertaining to the likelihood of peptide presentation calculated from each algorithm for the top scoring 10-percentile of n-mers categorized by HLA-allele (**F**) or n-mer length (**G**). **H.** Dot plot showing the overlay of the top scoring 10-percentile of both algorithms. Top-scoring final candidates are indicated in blue as the candidates that scored in the top 10 percentile in both algorithms. **I-J.** Bar graphs illustrating the composite presentation score (average presentation scores from HLAthena and MHCflurry 2.0) of top-scoring candidates binding across HLA-alleles based on frame-shift status (**I**) or alternative splicing category (**J**). **K-L.** Dot plot (**K**) and schematic (**L**) describing the Immunogenicity of the final candidate list based on top-scoring neojunction-derived n-mers derived from neojunctions detectable in patient-derived mass spectrometry and RNA sequencing data. **M.** Heatmap illustrating the intratumoral heterogeneity of final HLA-A*02:01-presented neojunctions across all spatially-mapped glioma samples. Glioma subtypes analyzed in this study include IDH1wt (blue), IDH1mut astrocytoma (yellow), IDH1mut oligodendroglioma (red), and one case (SF12548) that simultaneously exhibited either IDH1mut astrocytoma or IDH1mut oligodendroglioma across its spatially mapped samples (purple).

To test whether neojunctions are translated into proteins, we analyzed mass spectrometry (MS) data derived from glioma patients (*n*=447) in multiple publicly available MS data sets.^53–55^ Our analysis confirmed the detectable expression of unique neopeptides that map back to 302 (38.3%) unique public neojunctions **(Figure 4C).** We confirmed that these detected peptide sequences span the aberrantly spliced regions with sequence specific searches within the MS data and subsequent analysis of the resulting MS spectra **(Figures S4I-J)**. Interestingly, 41.7% of the detected peptides mapped back to neojunctions that encode for frame-shifts **(Figure S4K)**, signifying that frame-shift encoding splicing aberrations can still lead to detectable levels of translated peptides. From our peptidome analysis, we determined that neojunction-encoding transcripts are actively translated into detectable levels of protein products. When considering both RNA-seq and MS confirmation of glioma-specific neojunctions, we validated the presence of 192 (24.3%) public neojunctions expressed across all patient-derived samples **(Figure 4D)**, and thus, we looked towards identifying neoantigens derived from these verified neojunctions. Overall, these findings demonstrate the recurrence of public neojunctions across multiple transcriptomic and proteomic platforms.

### Tumor-wide neojunctions encode neoantigens predicted to be processed and presented by MHC-I

Reasoning that a fraction of neojunctions are translated and could be presented as targetable neoantigens^34,35,38^, we investigated whether the 789 characterized public neojunctions can generate peptides that are loaded onto MHC-I following proteasomal processing. All properly translated neojunction-derived sequences from TCGA were translated *in silico* to generate a neojunction-derived protein data set. Iterating through all possible n-mers of 8 to 11 amino acids **(Figure 4E)**, we defined tumor-specific n-mers as those absent from a UniProt reference normal human tissue proteome dataset.

Prediction of peptide presentation by MHC molecules based solely on binding affinity data often overlooks factors such as endogenous peptide processing steps.^56^ To more fully consider these features of neoantigens, we incorporated two independent prediction algorithms into our pipeline to identify neoepitope sequences that were likely to be processed and presented: MHCflurry 2.0 and HLAthena **(Figure S4A).** Each algorithm displayed robust capabilities in predicting either peptide processing motifs^57^ or pan-MHC binding affinities.^58^ The antigen processing predictor of MHCflurry 2.0 models allele-independent sequence properties and scored neoantigens favorably if their sequences (combined flanking amino acids and n-mer) are consistent with established motifs for efficient proteasomal cleavage.^57^ Meanwhile, the HLAthena’s prediction algorithm uses a large peptidome dataset comprised of 95 cell lines expressing single HLA class I alleles, allowing for independent HLA binding motif characterization.^58^ To rank the candidate public neopeptides, we assessed the binding potential of our n-mer candidates against the most prevalent human leukocyte antigen-A (HLA-A) haplotypes expressed across a wide range of demographics. As such, our pipeline investigated the presentation likelihood of neoantigen candidates by HLA-A*01:01, HLA-A*02:01, HLA-A*03:01, HLA-A*11:01, and HLA-A*24:02.^59^ To select high-binding targets, we focused on n-mer candidates that scored in the top 10% with both algorithms **(Figures 4F-G, S4B-S4C, Supplementary Table 2).** Candidate n-mers that yielded these scores (*n*=832) were predicted to be readily processed and presented in glioma and were retained for downstream analysis **(Figure 4H).** Mapping these top candidates to their originating neojunctions, we determined that 315 neopeptide-encoding neojunctions (NEJs; 39.92% of the originally characterized public neojunctions) produced cancer-specific peptide sequences containing top n-mer candidates. While a greater number of top-scoring n-mer candidates are generated from frameshifts and alternative exonic 3’ splice sites **(Figures S4D, S4E)**, presentation scores remained relatively consistent across n-mer candidates generated from NEJs with or without frameshift mutations or any particular alternative splice type **(Figures 4I-J, S4F, S4G).** To further narrow our NEJ candidate list, we cross-referenced the 315 NEJs with the 192 neojunctions that we previously characterized as being shared across transcriptomic and proteomic platforms **(Figure 4D).** Of the 192 neojunctions, 81 were characterized as NEJs through this analysis **(Figures 4K)**, with many NEJs encoding multiple strongly predicted candidates. We focused our downstream analyses on 32 NEJs that generate candidates that were predicted to bind strongly to HLA-A*02:01 due to the HLA allele’s high prevalence across North American and European populations^59,60^ and the ability to benchmark to other neoantigen studies^61–63^ **(Figure 4L)**. When ITH of these 32 neojunctions was investigated in the data set from spatially mapped samples, high intratumoral conservation was observed for most of these NEJs, particularly the neojunction located within *GNAS* (NJ*_GNAS_*) **(Figure 4M)**. These findings demonstrate that intratumorally conserved public neojunctions can potentially generate MHC-I-presented neopeptides that have potential for new immunotherapy development directed against tumor-wide targets.

### NEJ-reactive T-cell receptors can be isolated from donor CD8+ T-cells

We next sought to determine whether NEJ-derived neopeptides can drive T-cell immunogenicity. We performed *in vitro* sensitization (IVS) to stimulate and identify a neoantigen-reactive CD8^+^ T-cell population from healthy donor-derived peripheral mononuclear cells (PBMCs) using IFNg as a readout for reactivity^64,65^ **(Figure 5A)**. We focused our initial analysis on a subset (*n*=4) of our 32 top NEJ candidates that were predicted to generate high affinity binders to HLA-A*02:01 **(Figure 4K-4M)**. We therefore performed IVS of naïve CD8^+^ T-cells against neopeptide-pulsed autologous monocyte-derived dendritic cells (moDCs) collected from HLA-A*02:01^+^ healthy donors (*n*=5) with the goal of retrieving TCR gene sequences that confer specificity against these neoantigens. T-cells were co-cultured with mature dendritic cells pulsed with either NEJ-derived neopeptides, an influenza-derived peptide as a positive control, or no peptide as a negative control over the course of three 10-day cycles. To determine whether neoantigen-reactive CD8^+^ T-cells had expanded at the final timepoint, we cultured the sensitized CD8^+^ T-cells with T2 cells pulsed with neoantigen or influenza peptides. Because T2 cells are deficient in peptide transporters involved in antigen processing (TAP) proteins, they can be used to probe T-cell recognition of exogenous antigens in a non-competitive manner.^66,67^ Subsequent IFNγ ELISA assays on the corresponding T2:CD8^+^ conditions revealed neoantigen-reactive immunogenicity in two out of four of the public NEJ-derived neoantigens (NeoAs): NeoA_RPL22_ and NeoA_GNAS_ **(Figure 5B).** These results additionally indicate that NEJ-reactive CD8^+^ T-cells can exist within the naturally occurring human T-cell repertoire.

**Figure 5:**
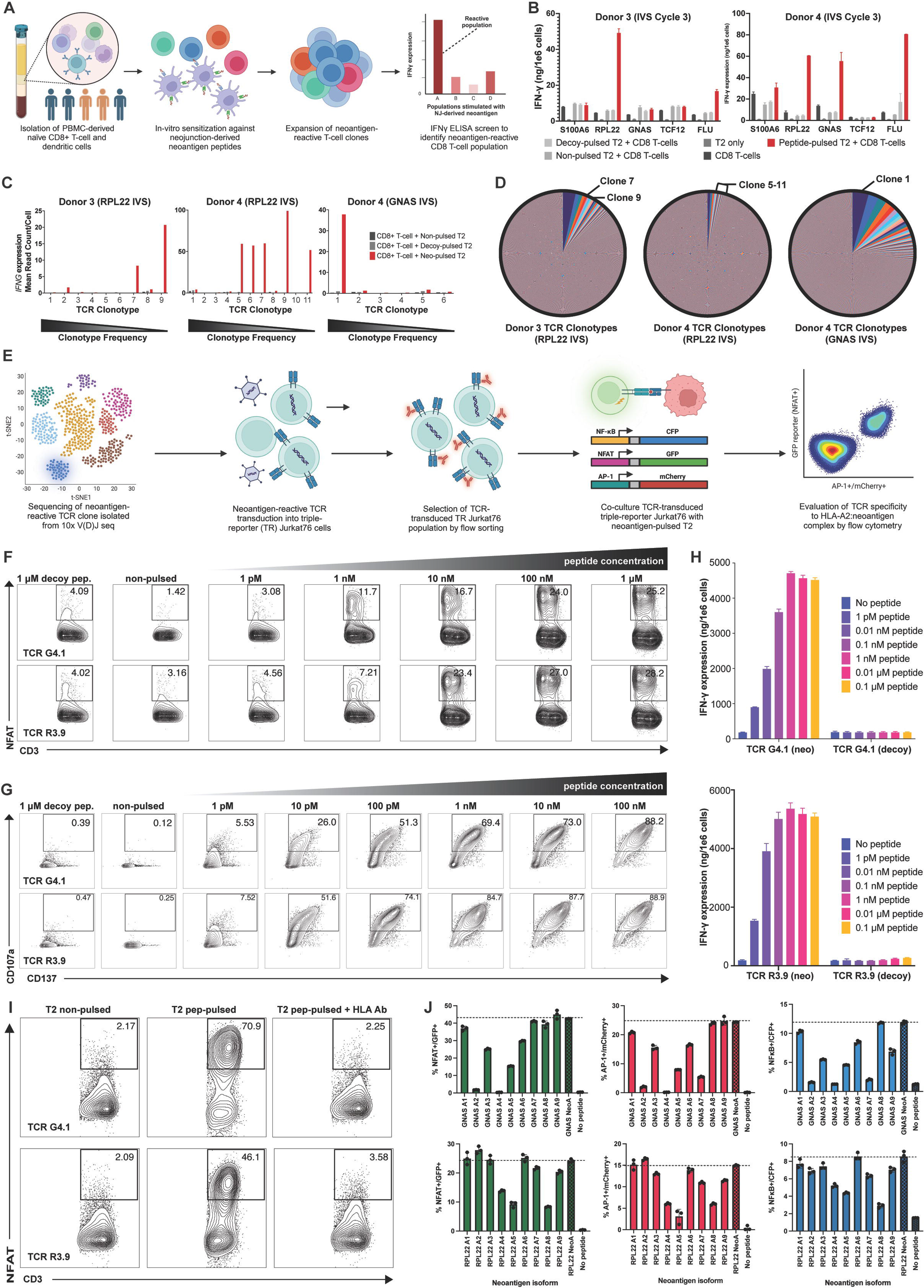
T-cell receptors (TCRs) specifically react to neojunction-derived neoantigens. **A.** Pipeline overview for identifying neojunction-derived neoantigen-reactive T-cell populations through *in vitro* sensitization (IVS) of healthy-donor PBMC-derived CD8^+^ T-cells against APC-presented neopeptides. **B.** IFNγ ELISA of reactive CD8^+^ T-cell populations following IVS with neoantigen. **C.** 10x V(D)J *IFNG* signatures of highly proliferated TCR clonotypes cultured against T2 cells pulsed with the neoantigen (colored), a control peptide (light-gray), or no peptide (dark-gray). Reactive TCR clonotypes were identified in Donor 3 CD8^+^ T-cells IVS-treated against NeoA_RPL22_ (left), Donor 4 CD8^+^ T-cells IVS-treated against NeoA_RPL22_ (center), and Donor 4 CD8^+^ T-cells IVS-treated against NeoA_GNAS_ (right). **D.** Clonotype frequency of all TCR clones identified in Donor 3 CD8^+^ T-cells IVS-treated against NeoA_RPL22_ (left), Donor 4 CD8^+^ T-cells IVS-treated against NeoA_RPL22_ (center), and Donor 4 CD8^+^ T-cells IVS-treated against NeoA_GNAS_ (right). **E.** Pipeline for validating the specificity of neoantigen-reactive TCR clonotypes found in 10x V(D)J single-cell RNA sequencing (scRNA-seq) against neojunction-derived neoantigen candidates utilizing a TCR-transduced triple-reporter Jurkat76/CD8 system followed by flow cytometry analysis. **F-G.** NeoA_RPL22_ (top) and NeoA_GNAS_-specific (bottom) TCR-transduced triple-reporter Jurkat76/CD8 cells (**F**) or PBMC-derived CD8^+^ T-cells (**G**) were activated against neoantigen-pulsed T2 cells in a dose-dependent manner. TCR-transduced cells were co-cultured with control peptide-pulsed T2 cells at the highest dose concentration (1mM). TCR activation of TCR-transduced triple-reporter Jurkat76/CD8 is measured by flow cytometry analysis of NFAT-GFP. PBMC-derived CD8^+^ T-cells were stained with CD107a and CD137 antibodies and surface expression of the TCR co-activation markers were analyzed by flow cytometry. **H.** IFNγ ELISA of NeoA_RPL22_ (top) and NeoA_GNAS_-reactive (bottom) TCR-transduced CD8^+^ T-cells co-cultured with dose-dependent neoantigen-pulsed (left) and control peptide-pulsed T2 cells (right). **I.** NeoA_RPL22_ (top) and NeoA_GNAS_-specific (bottom) TCR-transduced triple-reporter Jurkat76 cells are co-cultured with non-pulsed T2 cells (left), 0.1 μM neoantigen-pulsed T2 cells (center), or 0.1 μM neoantigen-pulsed T2 cells treated with HLA-A2 blocking antibody (right). Cells were stained with CD3 antibody, and TCR activation was evaluated by NFAT-GFP activity. **J.** Alanine scanning mutagenesis of NeoA_RPL22_ (top) and NeoA_GNAS_-reactive (bottom) TCR-transduced triple-reporter Jurkat76/CD8 cells co-cultured with alanine-substituted neoantigen-pulsed T2 cells, neoantigen-pulsed T2 cells, or non-pulsed T2 cells. Flow analysis was performed to evaluate TCR activity through NFAT-GFP (left), AP-1-mCherry (center), and NFκB-CFP (right) activity.

To retrieve the TCR gene sequences that confer reactivity to these neoantigens, we repeated the peptide-pulsed-T2:CD8^+^ T-cell co-culture assay on NeoA_RPL22_-and NeoA_GNAS_-reactive CD8^+^ T-cell populations and performed combined single-cell V(D)J and RNA-seq on all co-cultured conditions. As a quality control, we performed IFNγ and TNFα ELISA assays on supernatant from the same co-cultures generated for single-cell sequencing to confirm the presence or absence of neoantigen-reactive T-cell populations **(Figure S5A)**. Neoantigen-reactive TCR clonotypes were associated with significantly elevated *IFNG*, *TNFA*, *GZMB* transcripts when cultured with the corresponding neoantigen-pulsed APCs but not control peptide-pulsed or non-pulsed APCs.^65^ Using this method, we identified seven NeoA_RPL22_ -reactive TCRs, two from Donor 3 (TCR_R3.7_ and TCR_R3.9_) and five from Donor 4 (TCR _R4.5_, TCR _R4.6_, TCR _R4.7_, TCR _R4.9_, and TCR _R4.11_), and one NeoA_GNAS_-reactive TCR from Donor 4 (TCR_G4.1_) **(Figure 5C).** While only one NeoA_GNAS_-reactive TCR clonotype was characterized, this same clonotype was the most proliferated TCR clone, expanding to over 4% of the TCR repertoire in the CD8^+^ T-cell population **(Figure 5D).** The expansion of neoantigen-reactive CD8+ T-cell clones suggest a strong immunogenic proponent of these two neoantigens, with NeoA_GNAS_ demonstrating a much greater capacity in initiating T-cell expansion than NeoA_RPL22_. For the first time, we derived TCRs capable of recognizing cancer-specific neoantigens that are produced through recurrent mis-splicing events that are shared among patients and conserved across tumors.

### NEJ-reactive TCRs recognize NEJ-derived neoantigens in an MHC-I-restricted manner

To further test the specificity of the identified TCR_R3.9_ and TCR_G4.1_-reactive T-cell clones, we transduced TCR-null triple-reporter (TR) Jurkat76 cells which express the CD8a/b heterodimer (Jurkat76/CD8) or PBMC-derived CD8^+^ T-cells with lentiviral vectors encoding the retrieved TCR α- and β-chains. Jurkat76/CD8 cells do not express an endogenous TCR, allowing for the non-competitive introduction of exogenous TCRs.^67^ The TR Jurkat76/CD8 cells have response elements for NFAT, NF-κB, and AP-1 which drive expression of eGFP, CFP, and mCherry, respectively.^68,69^ Consequently, TCR activation can be quantified by measuring fluorophore expression levels **(Figure 5E)**. TCR-transduced TR Jurkat76 cells cultured with T2 cells pulsed with varying concentrations of neoantigen peptide demonstrated a dose-dependent reactivity **(Figures 5F, S5B, S5C).** Both TCRs demonstrated neoantigen-recognition against even low levels of peptide concentration (1 nM), illustrating high immunogenicity of the neoantigen target and high affinity of the corresponding TCR. The antigen-specificity of these receptors was supported by negligible TCR activation in the presence of supraphysiologic levels of the control peptide (1 μM). TCR-transduced PBMC-derived CD8^+^ T-cells further validated similar dose-dependent neoantigen-specific behavior **(Figure 5G, 5H).** TCR-transduced CD8^+^ T-cells were stained for surface expression of the T cell activation and degranulation markers, CD137 and CD107a, to quantify markers of T-cell activation and effector function. T-cell activation was observed at neoantigen-peptide pulsing concentrations as low as 1 pM **(Figure 5G).** Similarly, IFNγ and TNFα expression levels measured by ELISA suggested strong specificity by both TCRs as indicated by their EC_50_ between 0.01 to 0.1 nM **(Figure 5H, S5D).** To verify that T-cell activation is mediated by MHC-I-restricted presentation of the neoantigen, we treated neoantigen-pulsed T2 cells with an HLA-blocking antibody prior to co-culture with the TCR-transduced TR Jurkat76 cells. As expected, activation was completely blocked with the anti-HLA-I antibodies **(Figure 5I)**. Finally, we performed alanine scanning mutagenesis on both TCRs to determine whether there are off-target normal human proteins that share the peptide motif recognized by these TCRs.^70^ Single alanine mutations were sequentially introduced at each position within the NeoA_RPL22_ and NeoA_GNAS_ n-mer epitopes. TCR-transduced triple-reporter Jurkat76/CD8 cells were cultured against residue-substituted neoantigen isoforms, and key residues were defined by those that result in diminished TCR activation **(Figure 5J).** Alterations in recognition of a variant peptide indicates that the substituted residue was critical for TCR recognition.^71^ Referencing each TCR’s peptide recognition motif to a normal human proteome library (UniProt Proteome ID #UP000005640) demonstrated that there are no known human proteins that share the same set of key amino acid residues required for TCR recognition. This finding indicates that these TCRs specifically recognize the cancer-specific region of the neoantigen sequence. Taken together, our results reveal TCRs that exclusively recognize NEJ-derived public neoantigens with robust sensitivity and highlight a potential immunotherapeutic approach utilizing TCR-engineered T-cells to target this novel class of shared neoantigens.

### NEJ-derived public neoantigens are endogenously processed and presented by MHC-I

We next tested whether neojunction-derived transcripts are translated, processed by the proteasome, and presented at the cell surface by MHC-I. Presentation of NEJ-derived neoantigens was evaluated by two separate approaches: functional TCR-recognition and HLA-immunoprecipitation (HLA-IP) followed by subsequent liquid-chromatography-MS/MS (LC-MS/MS) **(Figure 6A).** While T2 cells can efficiently present exogenously introduced peptides, their deficiency for TAP proteins prevents them from performing endogenous antigen processing. Therefore, we transfected HLA-null COS7 cells with mRNAs encoding HLA-A*02:01 and the full-length neojunction-encoding transcript or the neoantigen peptide as a positive control.

**Figure 6:**
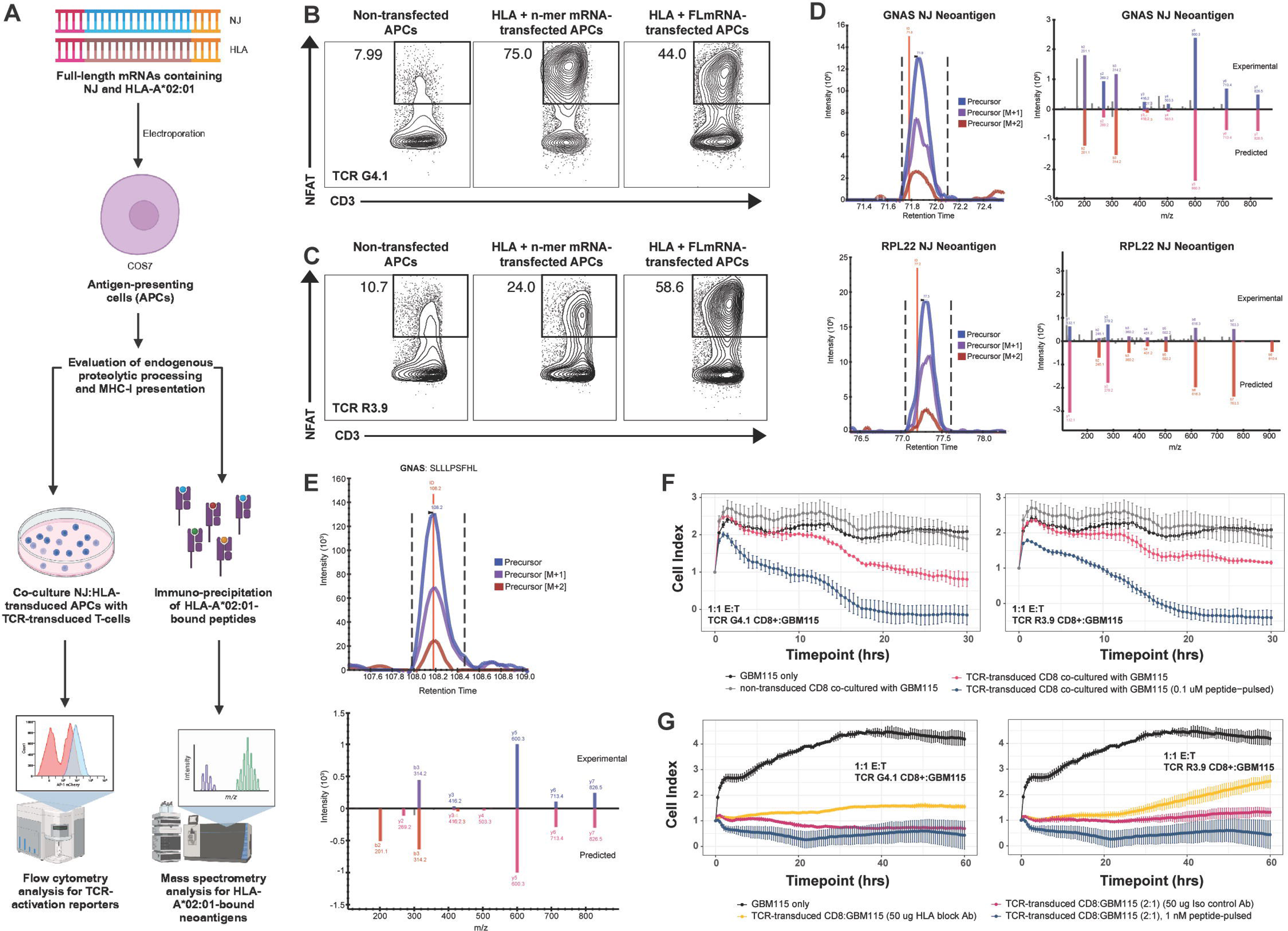
Neojunction-derived neoantigens are endogenously processed and presented by MHC-I to elicit neoantigen-specific TCR activation and tumor-specific killing. **A.** Pipeline overview for validating endogenous proteolytic cleavage of neoantigen candidates and the subsequent binding and presentation by MHC-I using two complementary methods. HLA-null antigen-presenting cells (APCs; COS7 or K562) were electroporated with mRNAs encoding the full-length gene with the neojunction mutation and HLA-A*02:01. First, TCR-activation following co-culture with APCs that express neojunction transcripts and HLA-A*02:01 was quantified by FACS using neoantigen-specific TCR-transduced triple-reporter Jurkat76/CD8 cells. Second, HLA-I-bound peptides were validated with IP-MS/MS of transfected APCs followed by identification of HLA-A*0201-bound peptide sequences. **B-C.** NJ_GNAS_-derived (**B**) and NJ_RPL22_-derived (**C**) neoantigen-specific TCR-transduced triple-reporter Jurkat76 cells were co-cultured against transfected COS7. Cultured cells were either non-transfected (left), transfected with mRNA encoding the neoantigen n-mer sequence and HLA-A*02:01 (center), or transfected with mRNA encoding the full-length (FL) mutant peptide and HLA-A*02:01 (right). TCR activation of TCR-transduced triple-reporter Jurkat76/CD8 was measured by flow cytometry analysis of NFAT-GFP. **D.** Mass spectrometry spectra of NJ_GNAS_-derived (bottom) and NJ_RPL22_-derived (top) neoantigen n-mers detected through IP-MS/MS following HLA-A*02:01 pulldown of HLA-A*02:01 and full-length neojunction-encoding mRNA-transduced COS7 cells. **E.** Mass spectrometry spectra of HLA-A2-presented NJ_GNAS_-derived neoantigen n-mers detected through IP-MS/MS following HLA-A*02:01 pulldown of GBM115 tumor cells. **F.** NJ_GNAS_-derived (left; colored), NJ_RPL22_-derived (right; colored) neoantigen-specific TCR-transduced, or non-transduced (gray) CD8^+^ T-cells were cultured against GBM115 tumor cells. Cell index indicates tumor cell adherence to the xCELLigence plate platform, with decreased cell index indicating tumor cell death. The assay was performed at an E:T ratio of 1:1. Cytotoxic killing was determined as the reduction of cell index compared to the control group with no CD8+ T-cell introduction (black) at a given timepoint. **G.** As controls, GBM115 tumor cells were incubated with an anti-HLA-I antibody (yellow), an isotype control antibody (purple), or pulsed with 1 nM of the neoantigen peptide (blue). NJ_GNAS_-derived (left) and NJ_RPL22_-derived (right) neoantigen-specific TCR-transduced CD8^+^ T-cells were cultured against these GBM115 tumor cells to test whether tumor killing is HLA-I dependent.

To demonstrate that expression of the neojunction at the transcript level can ultimately lead to a neoantigen-mediated immune response, we co-cultured COS7 cells transfected with the same HLA and full-length mutated transcript together with either TCR-transduced TR Jurkat76 or CD8+ T-cells. Similar to COS7 cells transfected with the mRNA encoding the neoantigen peptide, COS7 cells transfected with the full-length mutated transcript led to neoantigen-specific activation of both TCR_R3.9_ and TCR_G4.1_-transduced TR Jurkat76 and CD8+ T-cells, demonstrating that the identified public NEJs undergo all major biological steps required for proteolytic cleavage and MHC-I loading to enable T cell recognition (**Figure 6B-C**). Affinity-column-based immunopurification of HLA-I ligands was performed on HLA and mutant transcript-transfected COS7 cells. MS analysis showed that the same NeoA_GNAS_ peptide was identified as a high-confidence and high-abundance HLA-A2-bound peptide in HLA-A*02:01 and full-length NJ*_GNAS_*-transfected COS7 cells. Likewise, both NeoA_RPL22_ neopeptides were detected with high confidence in HLA-A*02:01, and NJ*_RPL22_*-transduced COS7 cells, with the higher scoring NeoA_RPL22_ 9-mer identified with higher relative abundance **(Figure 6D).** These direct experimental observations corroborate confirm our *in silico* predictions for proteasomal processing and HLA-binding **(Figure 4L).**

### TCR-transduced CD8^+^ T-cells mediate cytotoxicity against glioma cells expressing NEJ-derived public neoantigens

Based on the sensitivity of neoantigen-specific TCR **(Figures 5F, 5G),** we anticipated that endogenous levels of public NEJ expression in tumor cells would be sufficient to elicit a neoantigen-specific immune response. To test this hypothesis, we co-cultured the TCR_R3.9_- and TCR_G4.1_-transduced triple reporter Jurkat76/CD8 cells with IFNγ-pretreated HLA-A2^+^ LGG and GBM cell lines **(Figure S5E)** that were previously determined to express either NJ*_RPL22_* or NJ*_GNAS_* in RNA-seq **(Figure 4A)**. As expected, flow analysis following co-culture showed tumor-specific immune reactivity mounted by both TCR-transduced CD8^+^ T-cell lines against the glioma cell lines **(Figure 6E).** To evaluate whether neoantigen recognition is sufficient to induce a tumor-specific cytotoxic response, we transduced hybridized formats of these TCRs where the constant regions were replaced with a murine constant region to allow for accurate pairing of the transgene-encoded TCRa and TCRb hemichains. We then enriched for the transduced population of CD8+ T-cells through cell sorting using flow cytometry based on the positive staining of the murine TCR constant region. Tumor-specific cytotoxicity against endogenously-expressed NJ*_RPL22_* and NJ*_GNAS_* was assessed by co-culturing TCR-transduced CD8^+^ T-cells with the HLA-A*02:01^+^ glioma cell line GBM115. As a positive cytotoxicity control, we used neoantigen peptide-pulsed GBM115 cell lines to define maximum cell killing. At a 1:1 effector:target ratio, TCR_R3.9_ and TCR_G4.1_-transduced CD8^+^ T-cells efficiently demonstrated cytotoxic killing of 37.18% and 61.52%, respectively, of glioma cells, as detected using an impedance-based cell growth assay (xCELLigence) **(Figure 6F).** Increasing the effector:target ratio to 2:1 increased the potency of both clones, with 42.97% and 82.74% of tumor cells killed by TCR_R3.9_ and TCR_G4.1_-transduced CD8^+^ T-cells, respectively **(Figure 6F**; bottom). To confirm that tumor-specific killing is initiated by TCR recognition of the MHC:peptide complex, we introduced an HLA-I-blocking antibody into the co-culture conditions. The presence of anti-HLA-I antibodies partially blocked killing compared to an isotype control **(Figure 6G).** Here, we show that NEJs are endogenously processed and presented at sufficient levels in tumor cells for initiating tumor-specific cytotoxicity by neoantigen-specific CD8^+^ T-cells, highlighting a cell-based immunotherapy that targets a newly discovered repertoire of public, tumor-wide splicing-derived targets.

### Disease subtype-specific factors drive differences in neojunction expression

Noting the subtype-specific expression of neojunctions across IDH1wt, IDH1mut-A, and IDH1mut-O glioma subtypes in both TCGA and our spatially-mapped data set **(Figure 3E)**, we next sought to investigate dysregulation in splicing machinery that leads to upregulated neojunction expression. Both TCGA and our spatially-mapped RNA-seq data sets revealed that the total number of putative neojunctions per case is significantly greater in mutant *IDH1* gliomas compared to their wild-type counterparts **(Figures 7A, 7B).** Further dichotomies in neojunction expression were observed when *IDH1*wt, mutant *IDH1* astrocytoma (*IDH1*mut-A), and mutant *IDH1* oligodendroglioma (*IDH1*mut-O) glioma subtypes were compared. While *IDH1*mut-A gliomas demonstrated significantly higher average levels of neojunction expression compared to *IDH1*wt gliomas, *IDH1*mut-O neojunction expression far exceeds that of the other disease subtypes **(Figure 7C, 7D).** We next performed pairwise Pearson correlation analyses to explore whether neojunction expression is associated with somatic mutations in commonly mutated RNA splicing factors^72–74^ **(Figures S6A-S6C)**. We revealed high correlation values between *FUBP1*, *SF3A1*, or *NIPBL* with the *IDH1* mutation. *FUBP1* mutations have previously been reported as prevalent in *IDH1*mut-O glioma subtypes^75,76^. However, hierarchal clustering of neojunctions revealed no significant trend of neojunction expression with the mutation status of *FUBP1*, *SF3A1*, or *NIPBL* **(Figures S6D-S6I).** Prior studies have reported that dysregulation of individual splicing factors can result in aberrant splicing^73,77^. Based on these findings, we investigated whether aberrations in the expression levels of splicing-related genes correlates with the generation of neojunctions. To investigate possible drivers for the observed glioma-subtype differences in neojunction expression, we performed differential gene expression analysis (DESeq2) and gene set enrichment analysis (GSEA) on RNA-seq data from the three glioma subtypes from TCGA and looked for differentially expressed gene sets **(Figures S7A, S7B)**. GSEA highlighted significantly upregulated splicing-related gene sets in mutant *IDH1* cases compared to their wild-type counterpart across Gene Ontology Biological Processes (GOBP) **(Figure 7E)** and Gene Ontology Cellular Component (GOCC) databases **(Figure 7F).** When ordered based on increasing neojunction expression, splicing-related genes highly expressed in both mutant *IDH1* tumor subtypes largely clustered together **(Figure S7C).**

**Figure 7:**
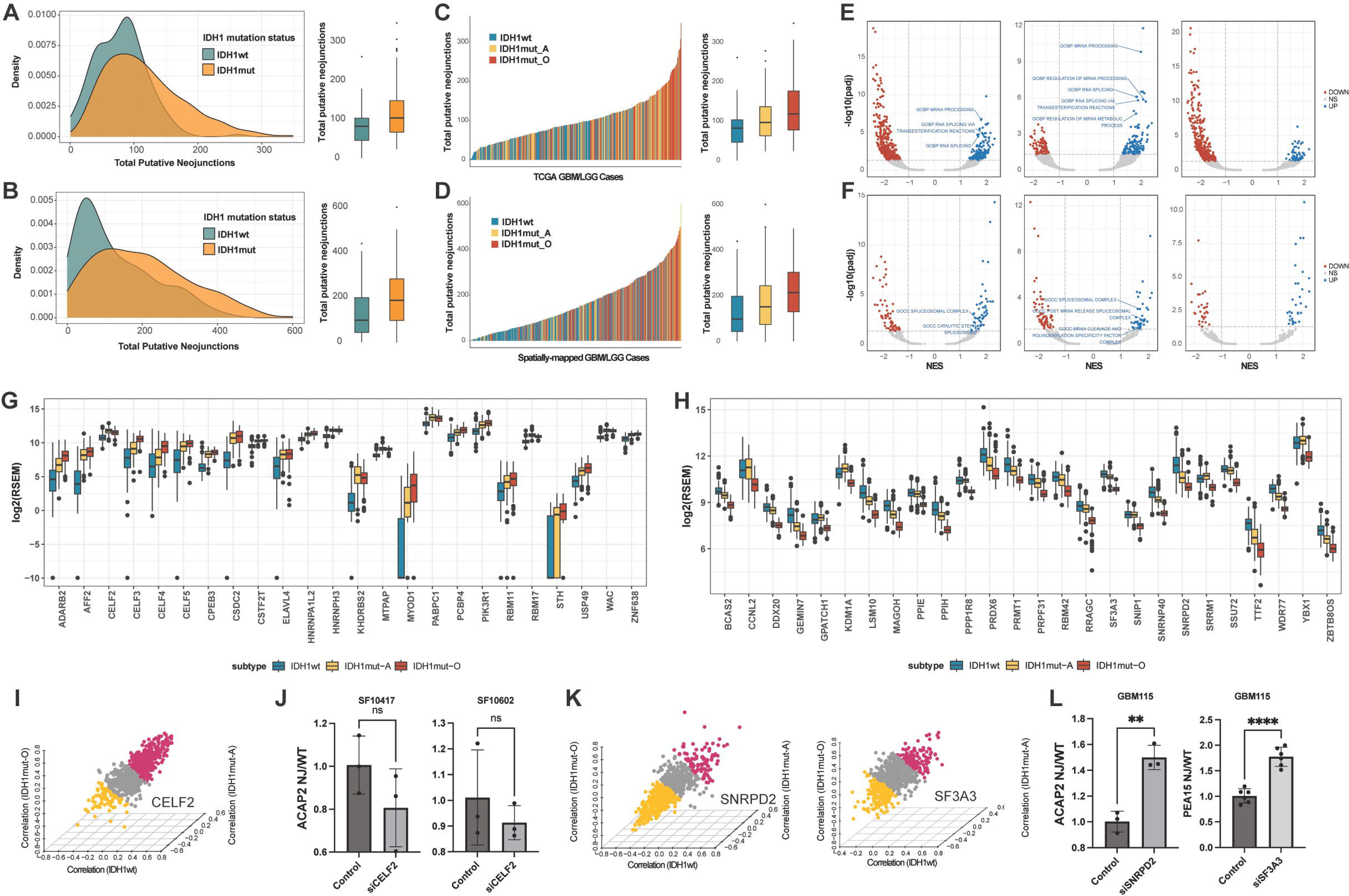
Glioma-specific disease subtypes demonstrate differential levels of neojunction expression. **A-B.** Density and box-and-whisker plots depict the total number of putative neojunctions expressed in IDH1mut cases (orange) and IDH1wt cases (green) in TCGA GBM/LGG samples (**A**) and our in-house spatially-mapped GBM/LGG dataset (**B**). **C-D.** Histogram and box-and-whisker plot depict the number total number of putative neojunctions expressed in *IDH1*wt (blue), astrocytoma (yellow), and oligodendroglioma cases (red) in TCGA GBM/LGG samples (**C**) and our in-house spatially mapped GBM/LGG dataset (**D**). **E-F.** Volcano plots illustrating significantly upregulated (blue) and downregulated (red) gene sets comparing *IDH1*mutO cases vs. *IDH1*wt cases (left), *IDH1*mutA cases vs. *IDH1*wt cases (center), and *IDH1*mutO cases vs. *IDH1*mutA cases (right). Gene sets categorized under the Gene Ontology Biological Processes (GOBP, **E**) and Gene Ontology Cellular Component (GOCC, **F**) were investigated in our analyses. Splicing-related gene sets were denoted by points labeled with text. **G.** Box-and-whisker plot depicting the log_2_(RSEM) expression level of splicing-related genes detected from GOBP gene set with a significant (*p* < 0.05) log_2_fold increase in expression of 1.5 between *IDH1*mutA (yellow) and *IDH1*mutO (blue) cases when compared to *IDH1*wt cases (red). **H.** Box-and-whisker plot depicting the log_2_(RSEM) expression level of chromosome 1p or chromosome 19q splicing-related genes detected from GOBP gene sets with a significant (*p* < 0.05) log_2_fold decrease in expression of 1.5 between *IDH1*mutO (blue) when compared to *IDH1*mutA (yellow) and *IDH1*wt cases (red). **I.** Pearson correlation analyses of glioma-specific neojunctions against the expression of *CELF2* in *IDH1*mutO (z-axis), *IDH1*mutA (y-axis), and *IDH1*wt (x-axis) cases. Neojunctions with a Pearson correlation greater than or equal to 0.10 with the corresponding gene are denoted with purple dots and those with a Pearson correlation less than or equal to - 0.10 with the corresponding gene are denoted with yellow dots. **J.** Expression of NJ_ACAP2_ in LGG cell lines, SF10417 (left) and SF10602 (right), treated with control siRNA or si*CELF2*. **K.** Pearson correlation analyses of glioma-specific neojunctions against the expression of *SNRPD2* (left) and *SF3A3* (right)in *IDH1*mutO (z-axis), *IDH1*mutA (y-axis), and *IDH1*wt (x-axis) cases.**L.** Expression of NJ_ACAP2_ in GBM115 treated with control siRNA or si*SNRPD2* (left) or si*SF3A3* (right).

To further investigate specific splicing-related genes that may lead to increased neojunction expression in mutant *IDH1* gliomas **(Figures 7G-H)**, we selected GOBP splicing-related genes (*n*=24) with a statistically significant (*p* < 0.05) 1.5-fold increase in expression in mutant *IDH1* cases compared to wild-type **(Figure 7G)**. Notably, *CELF2*^78,79^ and *ELAVL4*^80^ have previously been reported to generate splice aberrations when overexpressed. To this end, we employed correlation analyses of the expression of *CELF2* against the expression of all 789 public neojunctions identified. We identified a greater percentage of neojunctions whose expression generally increased (average Pearson correlation coefficient > 0.10) with the expression of both splicing-related genes across all glioma subtypes **(Figure 7I)**. Of the 789 neojunctions, 359 (45.5%) neojunctions increased in expression with *CELF2* expression as opposed to the 81 (10.3%) neojunctions that tended to decrease in expression with increasing levels of *CELF2*. Selecting the highest correlating neojunction associated with both splicing-related genes **(Figures S7D, S7E)**, we then performed siRNA-mediated knockdown of *CELF2* and *ELAVL4* in mutant *IDH1* cell lines, SF10417 and SF10602 **(Figure S7F)**, and observed a trend of decreased expression of the associated neojunction across both cases **(Figure 7J)**. These findings suggest that neojunction prevalence is mediated by tumor subtype-specific dysregulated expression of RNA-binding proteins and that modulation of these genes can effectively lead to changes in neojunction levels.

When revisiting GOBP splicing-related gene sets **(Figure S7C)**, subclusters of genes were significantly downregulated in *IDH1*mut-O cases. Most of the genes found within these clusters reside on either chromosome 1p or 19q. A distinctive diagnostic feature of *IDH1*mut-O gliomas is the co-deletion of chromosomes 1p and 19q. To evaluate whether the downregulation of these genes could lead to the characteristic increase in putative neojunction expression seen in the *IDH1*mut-O subtype, we selected GOBP splicing-related genes (*n*=26) with a statistically significant (*p* < 0.05) 1.5-fold decrease in expression in *IDH1*mut-O cases compared to both *IDH1*mut-A and *IDH1*wt cases **(Figure 7H).** Of these splicing genes, disruption of normal *SNRPD2* AND *SF3A3* expression was previously reported to lead to aberrations in splicing.^81^ Correlation analysis of *SNRPD2* and *SF3A3* expression against the expression of the 789 neojunctions across all glioma subtypes supports our hypothesis that decreased *SNRPD2* and *SF3A3* expression leads to greater neojunction expression **(Figure 7K)**. Of the 789 neojunctions, 385 (48.8%) neojunctions increased in expression with decreased *SNRPD2* expression compared with the 93 (11.8%) neojunctions that tended to increase in expression with increasing levels of *CELF2*. Similarly with increasing levels of *SF3A3* expression, 178 (22.6%) neojunctions tended to increase in expression and 127 (16.1%) neojunctions tended to decrease in expression. Notably, siRNA knockdown of *SNRPD2* and *SF3A3* in the GBM115 cell line **(Figure S7F)** led to a significant increase in the expression levels of their associated neojunctions **(Figure 7L).** This illustrates that the decreased expression of splicing-related genes due to the co-deletion of chromosomes 1p and 19q can lead to the upregulation of specific neojunctions. These findings suggest that commonly altered components of the RNA splicing machinery in gliomas are mechanistically linked to increased neojunction expression.

## DISCUSSION

Although several neoantigen-discovery pipelines have confirmed T-cell reactivity against tumor-specific neoantigens, none have considered their potential ITH, which is a major source of treatment failure. Furthermore, no study to date has investigated the ITH of neoantigens derived from RNA splicing aberrations. By incorporating intratumoral conservation of neoantigen expression in our computational pipeline, our workflow offers the advantage of addressing the challenges of ITH in treatment efficacy and ITH-related treatment failure. Here, we report on the discovery of novel class of intratumorally conserved public neojunctions resulting from dysregulated RNA splicing that generate tumor-specific peptides presented in the context of HLA-A*02:01, a highly prevalent HLA-I molecule. We further demonstrated that these neoantigens are immunogenic and their recognition by antigen-specific T cells can lead to tumor-specific killing. In validating the immunogenicity of NJ_GNAS_ and the specificity of TCR_G4.1_, we successfully identified a public tumor-wide NEJ that is robustly expressed intratumorally across multiple patients **(Figure 4N)**. When characterizing neoantigens, the extent of their presence throughout the tumor landscape is often overlooked, and in this study, we highlight a novel subset of NEJs that can be found tumor-wide through multiple biopsied sites. The endogenous processing and presentation of NeoA_RPL22_ and NeoA_GNAS_ further demonstrates that our workflow can accurately predict tumor-specific peptide candidates that are properly processed by the proteasome and strongly bind to MHC-I.

While we have investigated the immunogenicity of a subset of our predicted HLA-A*02:01-presented neoantigens, the NEJs with candidates that exhibited immunogenicity against specific TCR clones, NJ_RPL22_ and NJ_GNAS_, were identified in other cancers in our pan-cancer study as neojunctions, as well. Immunogenicity against specific TCR clones, NJ_RPL22_ and NJ_GNAS_, were identified as neojunctions in several other cancers we studied. Notably, our analysis of cohorts of multi-site samples indicate the expression of NJ_RPL22_ across multiple samples within the same tumor and the tumor-wide expression of NJ_GNAS_ in multiple tumor types **(Figure 3C).** The higher expression level of the canonical *GNAS* allele over *RPL22* may contribute to the prevalence of NJ_GNAS_ detected across all analyses **(Figure S3C).** This supports the observation of greater immunogenicity and tumor-specific killing by TCR_G4.1_ **(Figures 6F-H)** as there is a greater frequency in the generation and presentation NeoA_GNAS_ **(Figure 6E)**. Our findings indicate that the public NEJs identified in our study, notably NJ_GNAS_, can be targeted in cancer types beyond gliomas.

We also investigated the dysregulated expression of splicing-related genes associated with the *IDH1* mutation in gliomas because of the increased expression of neojunctions in these tumors. Mutations in *IDH1* and *IDH2* are prevalent in other cancers, including acute myeloid leukemia (AML)^82–84^, cholangiocarcinoma^85,86^, chondrosarcoma^87^, sinonasal undifferentiated carcinoma^88–90^, and angioimmunoblastic T cell lymphoma^91,92^. In our study, we successfully demonstrate that dysregulation in splicing factor expression can be seen in different disease types and that these aberrations can lead to significant changes in neojunction production. In the case of *IDH1* mutant oligodendrogliomas, *SNRPD2* expression is lost due to the tumor’s characteristic co-deletion of chromosomes 1p and 19q, and we demonstrate that targeted knockdown of this splicing-related gene leads to the direct increase in neojunction expression. This suggests that components of the RNA splicing machinery are mechanistically linked to the generation of neojunctions and targeting these components in future studies can potentially bolster targetable NEJ-derived neoantigen expression. Future analyses can establish whether similar splice-related genes are dysregulated in other cancers and whether their aberrant expression leads to a similar increase in neojunction levels. This question also may be relevant to mutations in splicing factors frequently found in tumors.^74^ While our analysis of mutant splicing factor *FUBP1* indicated modest changes in neojunction expression **(Figure S6C)**, we anticipate that splicing factor mutations in various cancers would lead to the generation of splicing anomalies.

### Limitations of the Study

The current study has focused on the identification of clonally conserved, immunogenic epitopes resulting from translated neojunctions that are restricted by prevalent MHC-I alleles. Several recent studies have identified HLA class II-restricted neoepitopes that are capable of driving antitumor responses in by neoantigen-reactive CD4^+^ T-cells in gliomas^93,94^ and other solid cancers^95,96^. Due to the limitations of currently available MHC-II binding prediction algorithms, our study did not assess whether HLA-II restricted public neoantigens resulting from clonally conserved NEJs are generated. CD4^+^ T-cells reactive against neoantigens may likely to be crucial for salvaging antitumor immune responses in cases where a tumor has lost MHC-I expression as an immune evasion mechanism. Furthermore, validation of neoantigen candidates from our pipeline was performed solely on predicted HLA-A*A02:01 binders to demonstrate a proof-of-concept. Future validation studies can be performed on the remaining candidates predicted as high-binders for other HLA alleles to further expand the repertoire of targetable neoantigens.

Importantly, the most comprehensive analysis of ITH (intratumorally-mapped samples ≥ 10) was conducted solely with GBM and LGG samples for this study due to the lack of prior studies that exceed 5 biopsies analyzed per tumor. As we previously mentioned, it is difficult to determine the number of samples required to fully recapitulate a tumor’s genomic landscape. In studies with only 3 intratumoral samples compared to our ≥ 10, the characterization of a neojunction as “tumor-wide” in our study holds greater statistical and biological weight. Thus, to fully validate a neojunction and its corresponding neoantigens as tumor-wide across other cancer types, future pan-cancer studies will likely need to include a greater library of intratumoral sites sequenced per sample and a wide anatomical distribution to maximally represent the tumor. Additionally, our study does not assess the biological contribution of the studied NEJs to the malignant phenotype. This can be relevant in understanding the clonal expression of tumor-wide NEJs and possible acquired resistance mechanisms.

Overall, our study highlights that RNA splicing aberrations are a robust source of intratumorally-conserved and tumor-wide public TSAs that the immune system can recognize. The ability to target these tumor-wide neoantigens with engineered T-cells enables a powerful therapeutic approach that could tackle the challenges of ITH that many neoantigen-based immunotherapies struggle with. Ultimately, the results from our study could allow us to design effective vaccine panels comprising tumor-wide neoantigen targets and engineer T-cell-based modalities that target tumor-wide splice-derived antigens across a wide range of cancer types.

## ACKNOLWEDGEMENTS

This study was supported, in part, by National Institutes of Health (NIH) grants R35NS105068 (H.O.), R01CA222965 (H.O.), NCI 2P50CA097257 (J.F.C), NCI P50CA097257 (J.F.C), NCI P01CA118816 (J.F.C), T32GM008568 (D.K.), 5T32CA151022 (L.C.), R37 CA259177 (C.A.K.), R01CA269733 (C.A.K.), R01 CA286507 (C.A.K.), P30 CA008748 (C.A.K.), NCI 1R50CA274229 (C. H.), Gianna Rae Meadows Grant for the Oligodendroglioma Cure (D.K.), Achievement Rewards for College Scientists Scholarship (D.K.), funding from the Glioblastoma Precision Medicine Project (J.F.C.), a generous gift from the Dabbiere Family, a generous gift from the Hana Jabsheh Research Initiative (J.F.C.), Brain Tumor Funders’ Collaborative (BTFC) (J.F.C and H.O.), the Parker Institute for Cancer Immunotherapy (H.O., C.A.K., and I.E.) and the Cancer Research Institute (I.E.). The UCSF Glioblastoma Precision Medicine Program is sponsored by the Sandler Foundation (J.F.C.).

## AUTHOR CONTRIBUTIONS

D.K. conceived the work and designed experimental setup and data analysis with input from H.O., J.C., and C.K. D.K and T.N. jointly designed and implemented the RNA-seq analysis pipeline and performed RNA-seq analyses, characterization of TCGA-derived neojunctions, and quantitative alternative splicing analysis. Pan-cancer analyses were the result of discussion among D.K., N.S., and J.C. D.K. analyzed publicly available MS data with guidance from A.B., E.R., and A.W. Spatially mapped samples and corresponding sequencing data were obtain through a collaboration between J.C., S.C., and J.P. D.K. and K.O. performed IVS against NEJ-derived neoantigens with input from P.W., H.O., A.S., and C.K. N.S. and C.H. provided help with 10x V(D)J scRNA-seq preparation, and L.C. assisted with scRNA-seq analysis and the identification of neoantigen-reactive TCR clones. D.K. performed all ELISA assays with assistance from G.C., A.D., and N.G. J.J. performed *in vitro* transcription to generate HLA and neojunction-encoding mRNA with experimental design input from I.E. Electroporation transcription and subsequent COS7:TR Jurkat76 co-cultures were performed by D.K. with input from I.E. Tumor killing assays using xCELLigence was conducted by D.K. with guidance from S.C. and A.Y. All flow cytometry data was obtained by D.K. with experimental setup assistance from A.Y and S.L. Flow cytometry analyses and gating was performed by M.G. and D.K. N.S. designed the primers for qPCR quantification of neojunction reads and performed siRNA KD experiments against splicing factors. D.K. wrote the manuscript, and all authors provided feedback on manuscript drafts.

---

**Supplementary Figure 1:**
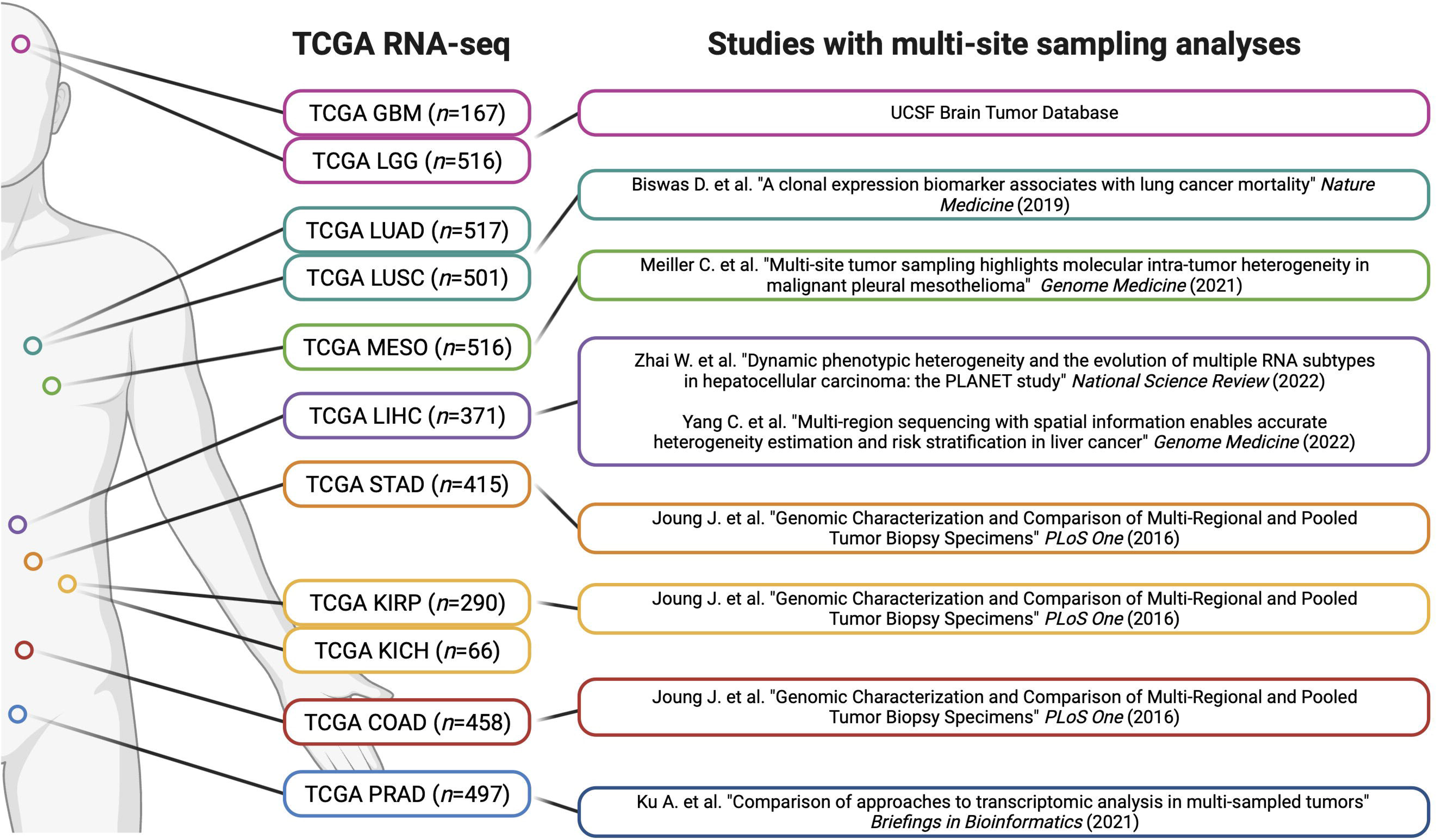
Multi-region intratumoral sampling datasets of various cancer types. Multi-region RNA-sequencing data of multiple cancer types were collected across various studies. Multi-region sampling is defined in studies in which multiple biopsies were isolated from same tumor for downstream sequencing analyses.

**Supplementary Figure 2:**
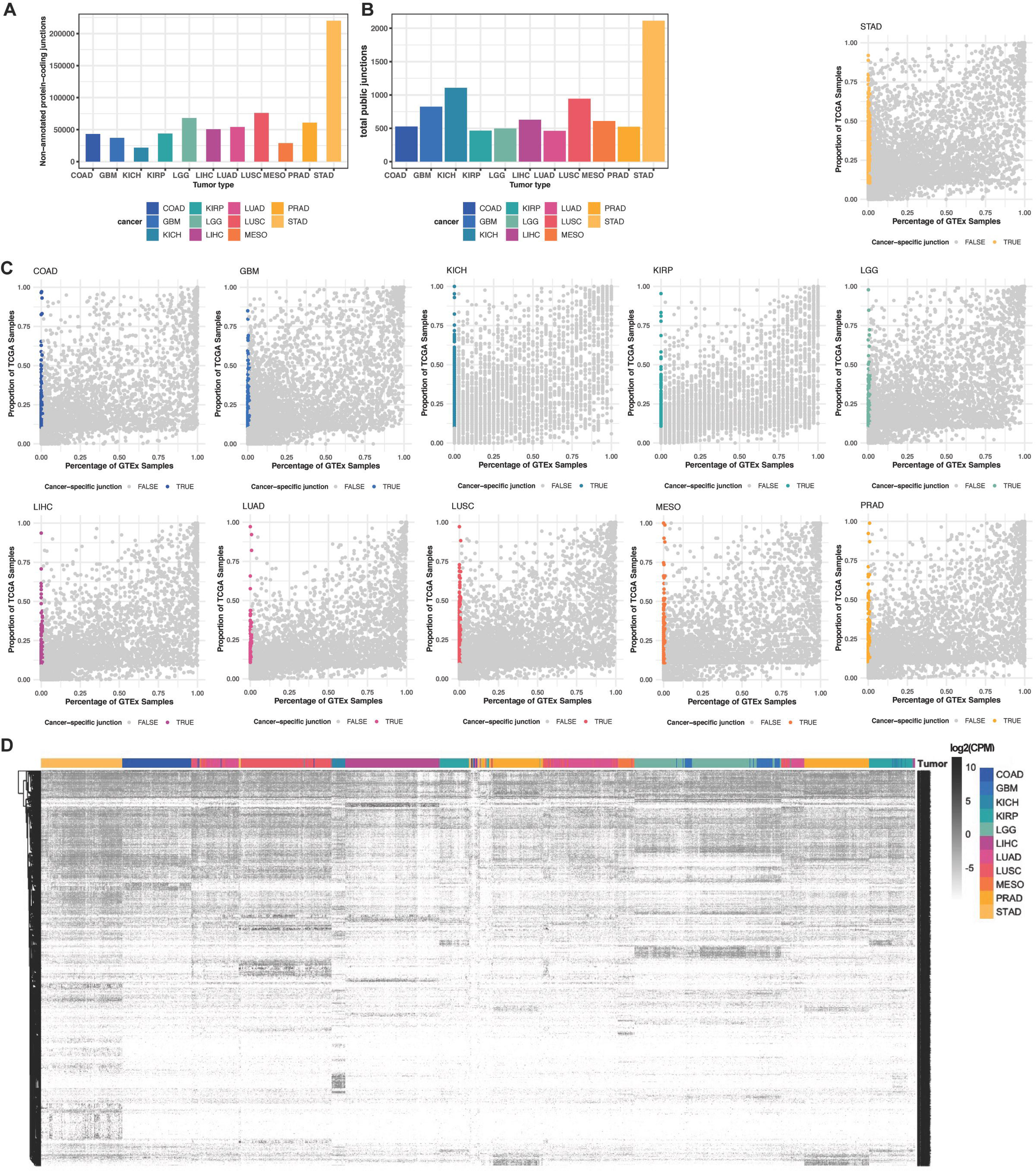
Pan-cancer public neojunctions are characterized from TCGA. **A.** Total number of non-annotated and protein-coding junctions detected pan-cancer. **B.** Total number of public (PSR_TCGA_ ≥ 10%), non-annotated, protein-coding junctions detected pan-cancer. **C.** Dot plots representing the positive sample rate percentage of non-annotated, protein-coding junctions in all studied cancer types (COAD, GBM, KICH, KIRP, LGG, LIHC, LUAD, LUSC, MESO, PRAD, STAD). Neojunctions (PSR_TCGA_ ≥ 10% and PSR_GTEx_ < 1%) are denoted by colored dots. **D.** Expression of all public neojunctions (log_2_(CPM)) that were detected across all studied TCGA tumor types. Unbiased hierarchical clustering was performed on all cases and neojunctions.

**Supplementary Figure 3:**
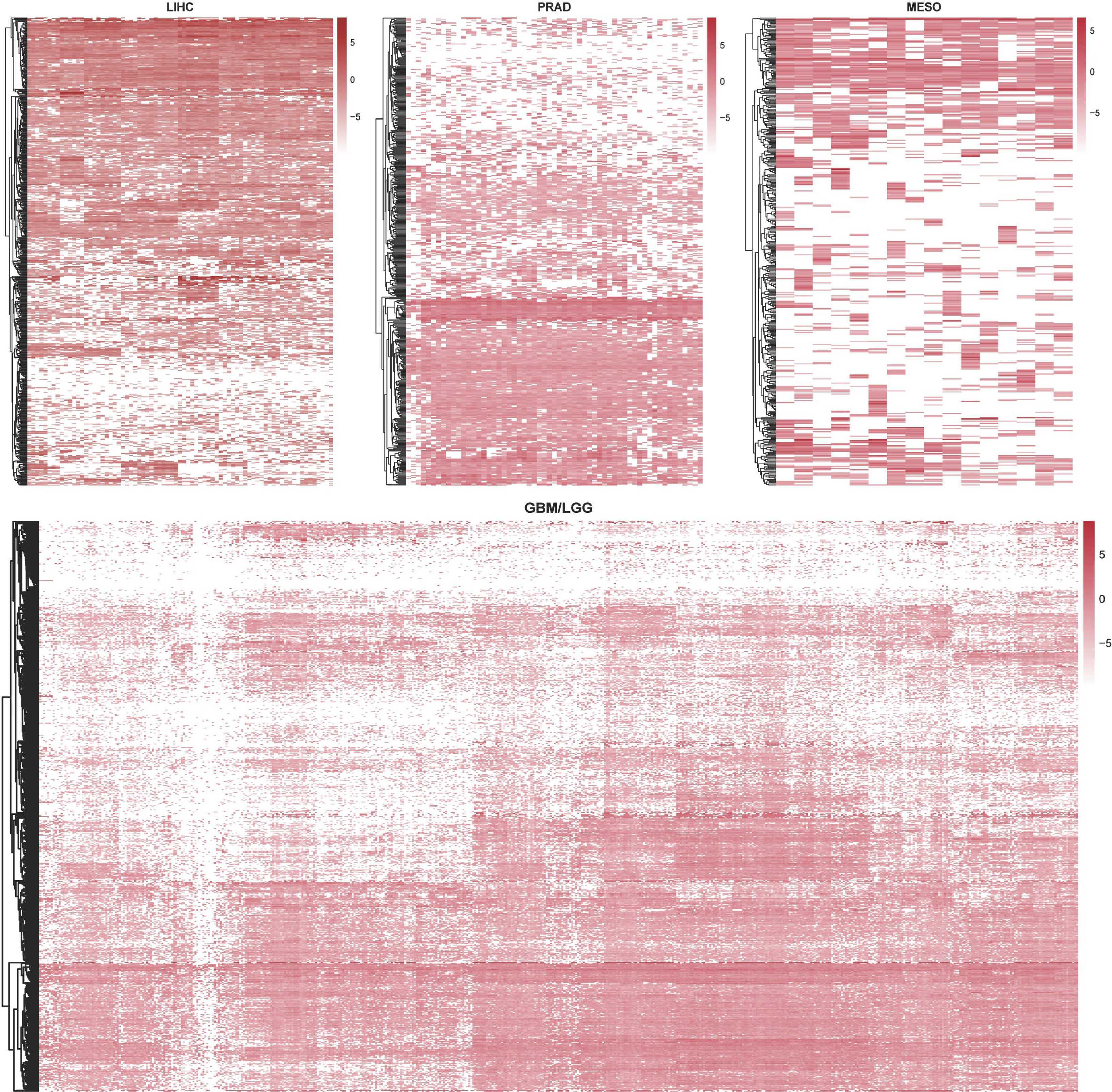
Intratumoral heterogeneity and interpatient characteristics of neojunctions across various cancer types. **A.** Counts per million (CPM) of non-annotated, protein-coding neojunctions across multi-region samples in **A.** kidney cancer **B.** prostate cancer, and **C.** mesothelioma, and **D.** glioma cases.

**Supplementary Figure 4:**
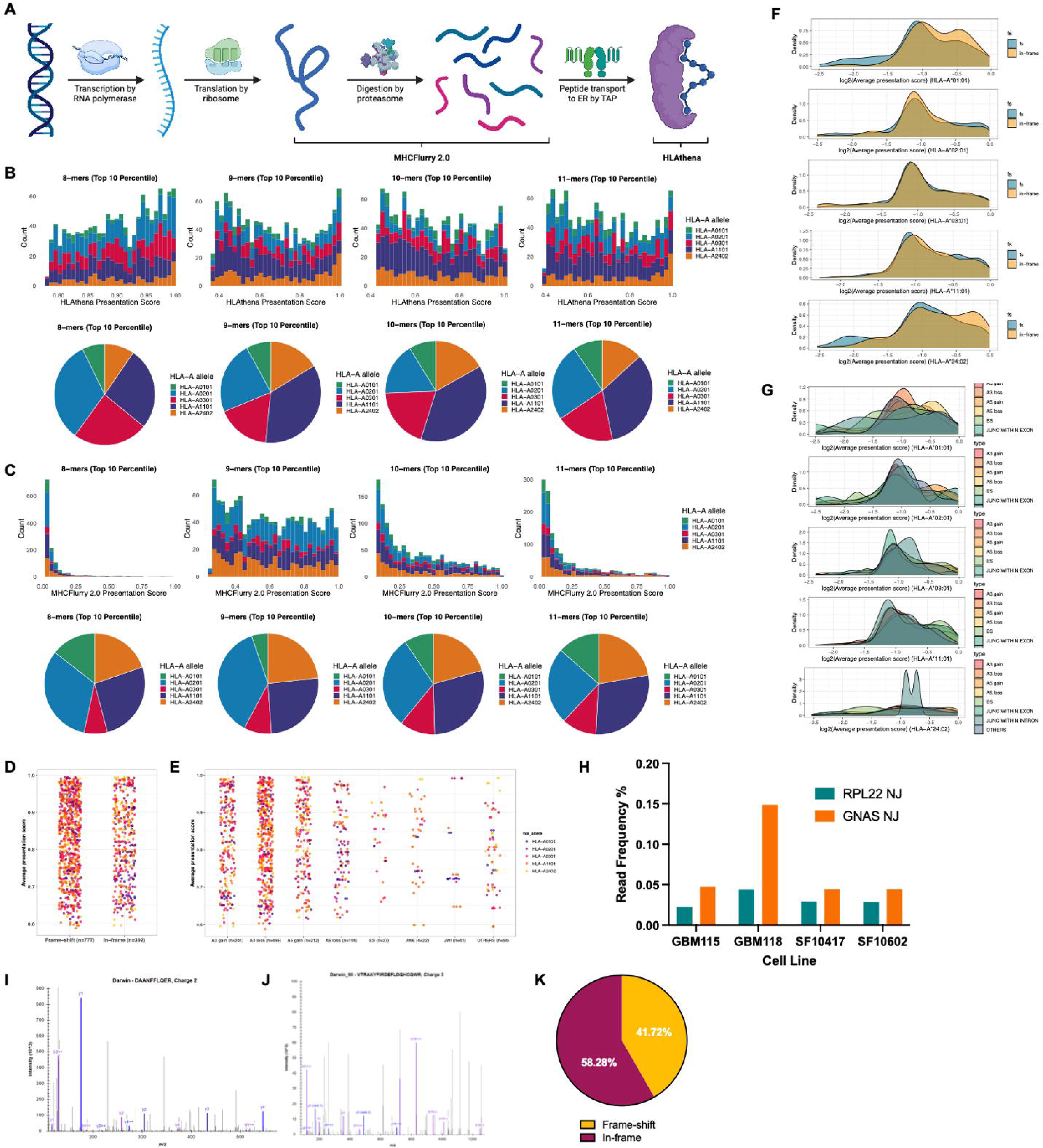
Neojunction-derived neopeptides are detectable in patient mass spectrometry and predicted to be processed and presented by MHC-I processed and presented by MHC-I. **A.** Schematic depicting the biological steps leading to the generation of MHC-I-presented antigens. Our neoantigen discovery pipeline considers the pre-presentation steps of proteasomal processing and HLA-binding. **B-C.** Histogram (top) and pie chart (bottom) depicting the distribution of the top scoring 8-mers, 9-mers, 10-mers, and 11-mers in **B.** HLAthena and **C.** MHCflurry 2.0. Scores pertain to presentation by HLA-A*01:01 (green), HLA-A*02:01 (blue), HLA-A*03:01 (red), HLA-A*11:01 (purple), and HLA-A*24:02 (orange). **D.** Jitter plot corresponding to the average presentation scores of peptides derived from neojunctions generating frame-shifts or in-frame mutations. **E.** Jitter plot corresponding to the average presentation scores of peptides derived from neojunctions derived from various splice types. **F-G.** Density plots depicting the average presentation scores of neoantigens derived from neojunctions generating **F.** frame shifts or **G.** various splice types presented by HLA-A*01:01, HLA-A*02:01, HLA-A*03:01, HLA-A*11:01, and HLA-A*24:02. **H.** Read frequency of reads spanning neojunctions in *RPL22* and *GNAS* compared to the canonical junction spanning reads in glioma cell lines (*n*=1). **I-J.** Mass spectra of peptide sequences spanning the aberrantly spliced regions in **I.** RPL22 and **J.** GNAS detected in publicly available GBM and LGG MS data. **K.** Proportion of mass spectrometry peptides that map back to neojunctions that encode for frame-shift or in-frame mutations.

**Supplementary Figure 5:**
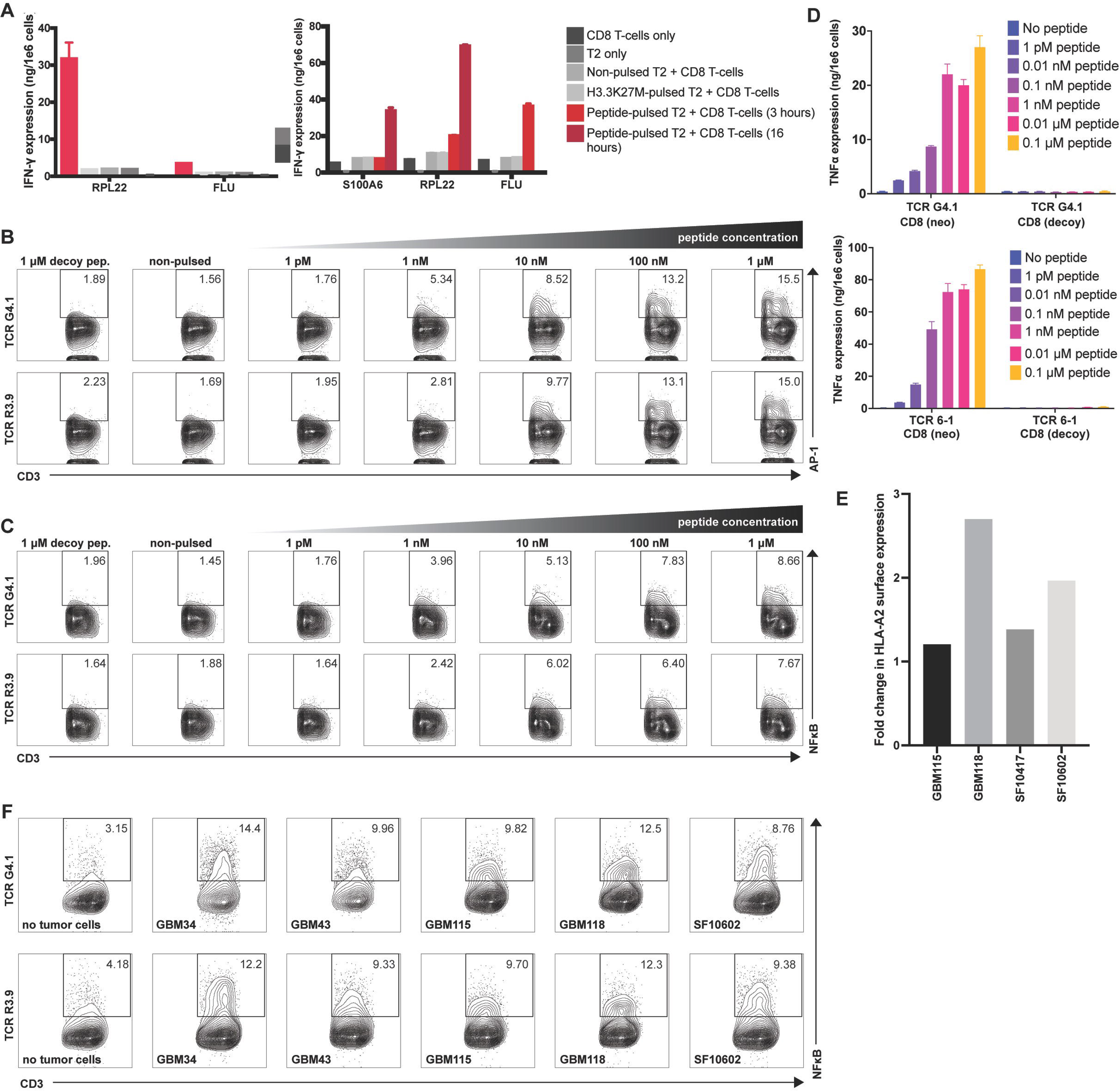
Neoantigen-reactive T-cell clones are isolated from PBMC and elicit an immune response upon neoantigen recognition. **A.** ELISA quantifying the expression of IFNγ in Donor 3 and Donor 4 in neoantigen-sensitized CD8+ T-cell populations (*n*=3) against T2 cells pulsed with the corresponding neoantigens. **B-C. B.** NJ_RPL22_-derived and **C.** NJ_GNAS_-derived neoantigen-specific TCR-transduced triple-reporter Jurkat76 cells activated against dose-dependent neoantigen-pulsed T2 cells. TCR activation of triple-reporter TCR-transduced triple-reporter Jurkat76 is measured by flow cytometry analysis of NFAT-GFP (top row) and NFκB-CFP (bottom row). **D.** TNFα expression of PBMC-derived CD8+ T-cells (*n*=3) transduced with TCR 6-1(top) and TCR 1-1 (bottom) against T2 cells pulsed with varying concentrations of corresponding neoantigen or decoy antigen. **E.** Surface HLA-A2 expression of LGG and GBM cell lines (*n*=1) with or without 48-hour pre-treatment of IFNγ.

**Supplementary Figure 6:**
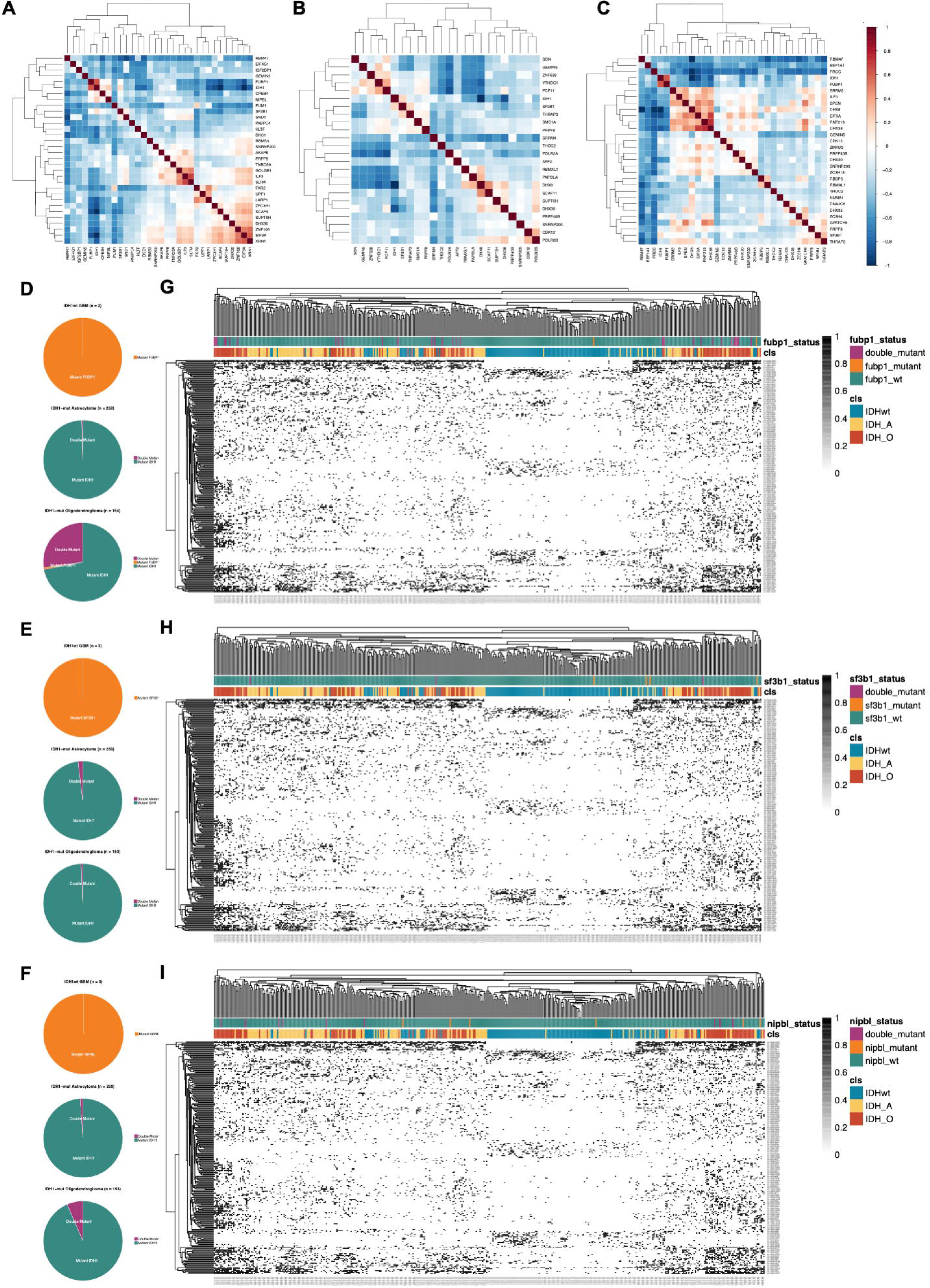
Co-occurrence of somatic mutations in splicing-related genes. **A-C.** Heatmaps showing the pairwise Pearson correlation matrix between gene expression of TCGA GBM/LGG samples computed for each gene pair. Splicing-related gene lists were defined by **A.** Nostrand et al. 2020, **B.** Sveen et al. 2016, and **C.** Seiler et al. 2018. **D-F.** Pie charts illustrating the proportions of IDH1 mutant samples that also contain mutations in **D.** FUBP1, **E.** SF3B1, and **F.** NIPBL in IDH1wt GBM samples (top), IDH1mut astrocytoma samples (middle), and IDH1mut oligodendroglioma samples (bottom). **G-I.** Binary heatmap demonstrating the putative expression of neojunctions in relation to glioma subtypes and mutation status of **G.** FUBP1, **H.** SF3B1, and **I.** NIPBL.

**Supplementary Figure 7:**
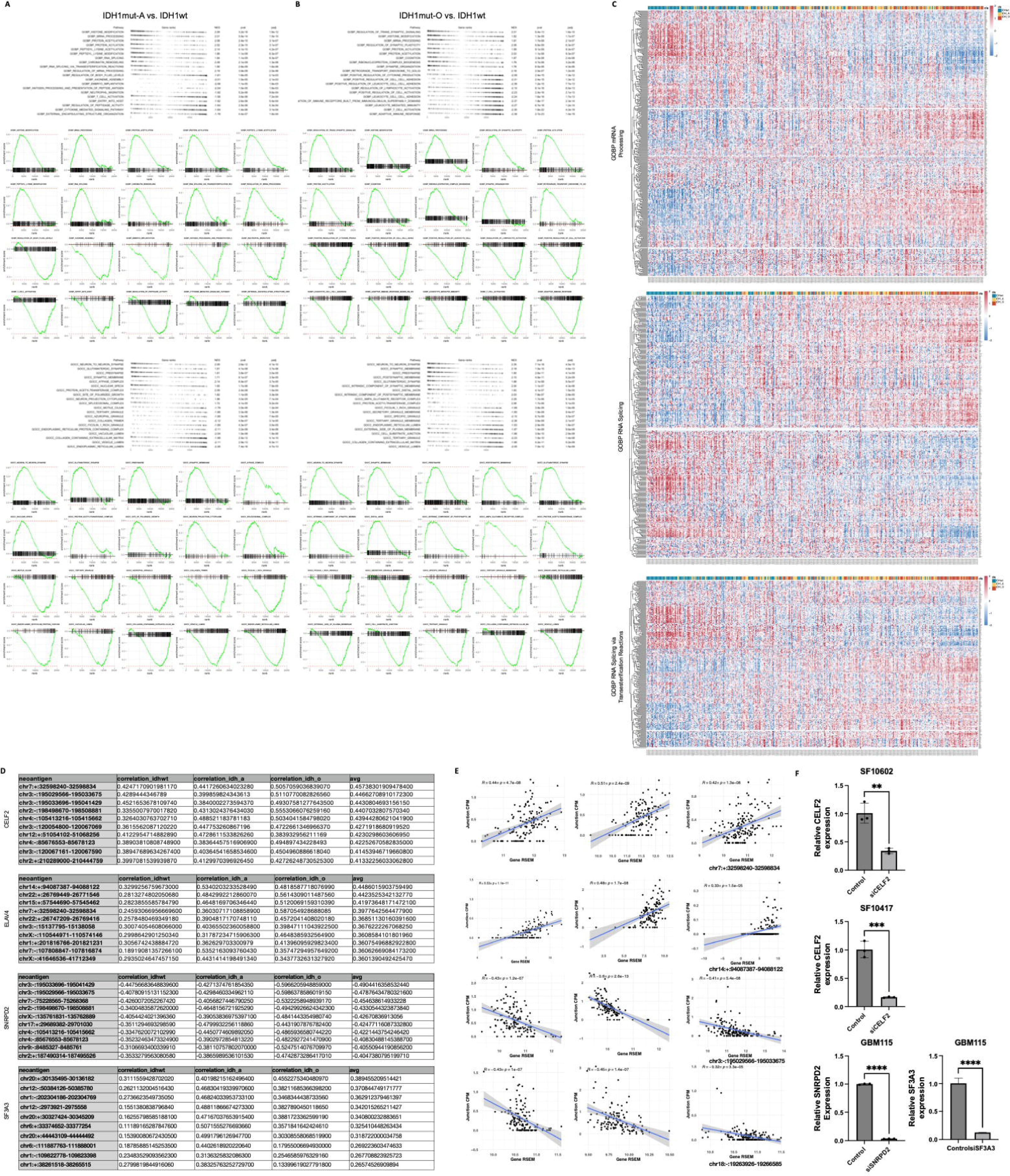
Glioma subtype-specific aberrations in splicing factor expression leads to differential levels of neojunction expression. **A.** Ranked log2 fold change of genes within the top enriched pathways within GOBP (top) and GOCC (bottom) when comparing IDH1mutA samples against IDH1wt samples. Green line plots show the running sum with the enrichment score peaking at the red dotted line, indicating leading edge genes that precede the peak. **B.** Ranked log2 fold change of genes within the top enriched pathways within GOBP (top) and GOCC (bottom) when comparing IDH1mutO samples against IDH1wt samples. **C.** Heatmap demonstrating hierarchal clustering of splicing-related genes (rows) in GOBP mRNA Processing (top), GOBP RNA Splicing (center), and GOBP RNA Splicing via Transesterification Reactions (bottom) in TCGA samples (columns) ordered by the total number of expressed putative neojunctions. **D.** Pearson correlation scores of the top 10 neojunctions that are positively correlated with *CELF2* and *ELAVL4* expression or negatively correlated with *SNRPD2* and *SF3A3* expression within all glioma disease subtypes. **E.** Pearson correlation dot plot of the top neojunction against the corresponding splicing-related gene. **F.** Expression levels of splicing-related genes in LGG and GBM cell lines treated with siRNA. SF10602 (top) and SF10417 (center) were treated with siRNA targeting *CELF2*. GBM115 was treated with siRNA targeting *SNRPD2* (bottom-left) and *SF3A3* (bottom-right)

## RESOUCE AVAILABILITY

### Lead contact

Further information and requests for resources and reagents should be directed to and will be fulfilled by the lead contact, Hideho Okada.

### Materials availability

This study did not generate new unique reagents.

### Data and code availability

Spatially-mapped biopsy RNA sequencing data will be deposited. HLA-IP and LC-MS/MS data will be deposited and made publicly available at the date of publication. Additionally, single-cell V(D)J sequencing data of identified TCRs will be deposited and made publicly available as of the date of publication. Accession numbers are listed in the key resources table. All original code for the identification of tumor-wide public neojunctions have been deposited at GitHub (https://github.com/dakwok/SSNIP) and are publicly available as of the date of publication. Any additional information required to reanalyze the data reported in this paper is available from the lead contacts upon request.

## EXPERIMENTAL MODEL AND STUDY PARTICIPANT DETAILS

### Human clinical datasets

The intratumoral multi-region sampling cohort for various cancer types utilizes RNA sequencing data from the following studies:

1. This paper, for multi-region sampling of glioblastoma and low-grade glioma
2. Yang et al. (Genome Medicine, 2022), for multi-region sampling of hepatocellular carcinoma.
3. Joung et al. (PLoS One, 2016), for multi-region sampling of hepatocellular carcinoma, stomach adenocarcinoma, renal cell carcinoma, and colon adenocarcinoma.
4. Ku et al. (Briefings in Bioinformatics, 2021), for multi-region sampling of prostate cancer.
5. Meiller et al. (Genome Medicine, 2021), for multi-region sampling of mesothelioma.
6. Bakir et al. (Nature, 2023), for multi-region sampling of non-small cell lung cancer

Analysis of neojunction expression within multi-region samples were conducted immediately with our neojunction prediction pipeline if the FASTQ file is available. If RNA-sequencing data is only available in BAM format, the sequencing file is converted into FASTQ format utilizing the Picard software (version 2.7.7a). Neojunction prediction is detailed in the Method Details section.

## METHOD DETAILS

### Data download

Bulk RNA-sequencing data for glioblastoma (GBM; n=167), low-grade glioma (LGG; n=516), lung adenocarcinoma (LUAD, n=517), lung squamous cell carcinoma (LUSC, n=501), mesothelioma (MESO, n=516), liver hepatocellular carcinoma (LIHC, n=371), stomach adenocarcinoma (STAD, n=415), kidney renal clear cell carcinoma (KIRC; n=533), kidney renal papillary cell carcinoma (KIRP; n=290), kidney chromophobe (KICH, n=66), colon adenocarcinoma (COAD; n=458), and prostate adenocarcinoma (PRAD; n=497) samples were downloaded from TCGA in FASTQ format. Download of intratumoral multi-region sampling sequencing data is detailed in the previous section. Similarly, bulk RNA-sequencing data for 9,166 normal tissue samples in FASTQ format were downloaded from the Genotype-Tissue Expression (GTEx) repository. Bulk-RNA sequencing data for 66 patient-derived GBM cell lines were received from the Mayo Clinic Brain Tumor Patient-Derived Xenograft National Resource.^1^ Proteomics data for 100 GBM samples were downloaded from the Clinical Proteomic Tumor Analysis Consortium (CPTAC).^2^

### RNA sequencing alignment

All downloaded RNA-sequencing data sets were individually aligned using a STAR aligner-based processing pipeline. Using the STAR software (version 2.7.7a), we constructed a genome index containing non-annotated junctions through the initial alignment pass of the input data. The complete set of command line parameters: --runThreadN 1 \ -- outFilterMultimapScoreRange 1 \ --outFilterMultimapNmax 20 \ --outFilterMismatchNmax 10 \ --alignIntronMax 500000 \ --alignMatesGapMax 1000000 \ --sjdbScore 2 \ --alignSJDBoverhangMin 1 \ --genomeLoad NoSharedMemory \ -- limitBAMsortRAM 80000000000 \ --readFilesCommand gunzip -c \ --outFilterMatchNminOverLread 0.33 \ -- outFilterScoreMinOverLread 0.33 \ --sjdbOverhang 100 \ --outSAMstrandField intronMotif \ --outSAMattributes NH HI NM MD AS XS \ --limitSjdbInsertNsj 2000000 \ --outSAMunmapped None \ --outSAMtype BAM SortedByCoordinate \ --outSAMheaderHD @HD VN1.4 \ --twopassMode Basic \ --outSAMmultNmax 1 \ and aligned using the GRCH37 STAR index file.

### TCGA sample selection and gene expression quantification

TCGA tumor samples with an absolute tumor purity greater than 0.60 were retained downstream *in silico* analysis. (Aran et al. 2015, Ceccarelli et al. 2016) We selected non-mitochondrial, protein-coding transcripts defined by the Ensembl Homo Sapiens GRCH37.87 gene annotation gene transfer format (GTF) file and utilized this curated to select and retain protein-coding transcript isoforms within the TCGA RNA-sequencing data. Transcript-level expression data (log2[RSEM-TPM+0.001]) for all TCGA samples were downloaded from the UCSC Xena Toil-pipeline and transformed into standard TPM values. Protein-coding transcript isoforms with a median TPM ≥ 10 were retained for downstream analysis. In the case of glioma TCGA cases, subsequent expression data in TPM was subset into 6 disease type categories: all cases (*n*=429), GBM cases (*n*=115), LGG cases (*n*=314), *IDH1*-WT cases (*n*=166), *IDH1*-MUT astrocytoma cases (*n*=140), *IDH1*-MUT oligodendroglioma (*n*=123). Protein-coding transcript isoforms with a median TPM ≥ 10 in at least one of the 6 disease types were retained for further analysis.

### Characterization of public neojunctions

For public cancer-specific splicing event counting, we designed a custom R script that detected and quantified non-annotated, cancer-specific splicing events found across each corresponding patient cohort. From the output files derived from STAR aligner in the previous step, alternative splicing events were quantified in detected junction counts within the corresponding sj.out.tab file. We removed splicing events detected in the GRCh37.87 GTF sj.out.tab (GENCODE v33) file to define non-annotated splicing junctions. Non-annotated splicing junctions that overlap non-mitochondrial, protein-coding genes identified in the previous step were retained for continued analytical processing. We removed all splicing junctions with less than 10 of its target spliced reads (count) or less than 20 total spliced reads (depth) over the whole cohort. Similarly to previous studies^3^, we computed spliced frequency as the sum of the total number of target spliced reads divided by the collective sum of spliced reads from the target and canonical junctions. Splicing junctions with a read frequency greater than 1% were retained for downstream analyses. We defined public splicing junctions as ones that were putatively expressed with the aforementioned criteria of total read count, read depth, and read frequency across at least 10% of the studied patient cohort and retained those for further analysis. To characterize cancer-specific splicing events, otherwise known as neojunctions, we removed all junctions that putatively expressed with the same parameters in more than 1% of GTEx normal samples.

### Detection of cancer-specific intron retention events

Intronic splicing events were detected and characterized using IRFinder v1.2.3. RNA sequencing data from TCGA (GBM/LGG) and GTEx (CNS) aligned to GRCh37 (hg19) were imported into the software for the detection of intron retention events. General linear model (GLM)-based analysis was used for differential intron retention assessment. The intron retention ratio is calculated as (intronic reads/sum(intronic reads, normal spliced reads). Significant intron retention changes are defined as (1) no less than 10% in both directions (2) adjusted p-values less than 0.05. An intron retention event’s PSR within TCGA or GTEx is defined as the number of cases that fulfill these criteria divided by the total number of cases within the cohort. Putative cancer-specific intron retention neojunctions are characterized as intron retention events with a TCGA PSR ≥ 0.10 and a GTEx PSR < 0.01.

### Transcriptomic validation of expressed neojunctions

<u>Detection of expressed neojunctions in patient-derived GBM/LGG cell lines:</u> RNA sequencing data derived from GBM PDX cell lines were downloaded from the Mayo Clinic Brain Tumor Patient-Derived Xenograft National Resource. Patient-derived LGG cell lines were generated from surgically-resected specimens in University of California, San Francisco’s Neurological Surgery Brain Tumor Center.^4^ RNA sequencing data from GBM and LGG cell lines were aligned and processed as described above. Public neojunctions with splice junction counts per million (CPM) > 0 are considered detectable in cell line-derived RNA sequencing data. <u>Detection of expressed neojunctions in multi-region</u> <u>cases:</u> In our cohort of spatially-mapped glioma cases, approximately ten or more maximally-distanced anatomical biopsies were collected from each patient, allowing for intratumoral assessment of genetic heterogeneity via bulk RNA-seq and whole-exome sequencing. Multi-region sequencing data of various other cancer types vary in the number of sampled regions per tumor and are detailed in the corresponding references **(Figure S1)**. RNA-sequencing data collected from each multi-region sample was processed and aligned as described above. We searched for putative neojunctions previously characterized from TCGA within each multi-region sampling dataset. Public neojunctions with CPM > 0 were considered detectable. Public neojunctions with putative expression (≥ 10 spliced reads) in two or more mapped samples within the same case are considered spatial-conserved neojunctions. Neojunctions detected in all multi-region samples within the same tumor are considered tumor-wide neojunctions.

### Proteomic validation of expressed neojunction-derived peptides

From the putative neojunctions detected in the above pipeline, we generated a database of all plausible polypeptides derived from all neojunctions. Neojunction-encoding transcripts were generated by mapping the junction coordinates to an hg19 human genome assembly within the Ensembl annotation database (AH13964, EnsDb.Hsapiens.v75). Prediction of neojunction-derived amino acid sequences were subsequently performed, and appropriately translated sequences (methionine starting residue, removal of sequences following first stop codon) were retained for downstream n-mer iteration. To detect neojunction-derived polypeptides within GBM cases, we analyzed .RAW files of GBM and LGG MS dta housed in the Clinical Proteomics Tumor Analysis Consortium (CPTAC, *n*=99), Bader et al. (*n*=99), Lam et al. (*n*=92), and Yanovich-Arad et al. (*n*=84). MaxQuant (v1.6.17.0) was used to identify tryptic sequences from the corresponding MS data sets. Predicted neojunction-derived peptides, decoy sequences, and a human reference proteome (UniProt Proteome ID #UP000005640) were inputted as a FASTA file into MaxQuant, and tryptic sequences derived from the input file were matched against the publicly available MS databases. Cancer-specific peptides spanning neojunction-derived protein sequences were considered MS-confirmed. The relative detection levels of the neojunction-derived peptides and normal tissue-derived peptides were evaluated by their log2(peak intensities). Aside from the default settings, the following commands and parameters were modified and used for MS analysis in MaxQuant: Digestion mode = Trypsin/P; Max missed = 3; Minimum peptide length = 5; Minimum peptide length for unspecific search = 5.

### Peptide processing and MHC-I binding and presentation predictions

Cancer-specific transcripts with associated neojunctions were translated *in silico* into their corresponding amino acid sequences. A library of all possible peptides of 8 to 11 amino acids in length was then generated, and cancer-specific sequences were selected by removing those detectable in normal tissue peptide isoforms in a reference human proteome dataset (UniProt Proteome ID #UP000005640). All cancer-specific peptides with their upstream and downstream flanking sequences (maximum flanking length of 30 amino acids) were independently analyzed and ranked by MHCFlurry 2.0 and HLAthena MSiC. HLA-I binding affinity was assessed against HLA-A*01:01, HLA-A*02:01, HLA-A*03:01, HLA-A*11:01, and HLA-A*24:02 in both cases. In the HLAthena evaluation of antigen binding and presentation to the corresponding HLA haplotypes, peptides were assigned to alleles by rank with a threshold of 0.1. Context of up to 30 flanking amino acids on both N and C terminus were utilized with aggregation by peptide and no log-transformed expression. Baseline MHCFlurry 2.0 models with both peptide:MHC-I binding affinity (BA) predictor and antigen processing (AP) predictor was used. Overall, peptide:HLA presentation scores were characterized by mhcflurry_presentation_score and MSiC_HLA scores in MHCFlurry 2.0 and HLAthena, respectively. To select for high-binders, we curated lists of peptide:HLA complexes within the top 10 percentile of scores from both prediction algorithms.

### Cell culture

<u>GBM PDX cell culture:</u> Patient-derived xenograft (PDX) glioblastoma cell lines, GBM34, GBM43, GBM108, GBM115, GBM118, GBM102, GBM137, GBM148, GBM164, and GBM195, were obtained from the Mayo Clinic Brain Tumor PDX national resource. Xenograft lines were cultured as by recommended conditions in previous literature^5,6^ and passaged a maximum of 20 times before restoration to earlier passages. Cells were cultured in Dulbecco’s Modified Eagle’s medium (DMEM) supplemented with 10% fetal bovine serum and 1% penicillin and streptomycin (P/S). Cell culture plates were treated overnight at 4 °C with DPBS (with calcium and magnesium) and 10% laminin (Gibco^TM^ Cat. #23017015) prior to use. <u>Primary patient-derived GBM/LGG cell culture:</u> Primary patient-derived wildtype *IDH1* GBM (SF7996), mutant *IDH1* astrocytoma (SF10602), and mutant *IDH1* oligodendroglioma (SF10417) cell lines were previously internally generated from dissociated glioma biopsies and cultured as previously described.^4^ Cells were cultured in serum-free, glioma neural stem (GNS) cell medium, which comprises of Neurocult NS-A (STEMCELL Technologies Cat. #05751) supplemented with N-2 supplement (Invitrogen Cat. #17502048), B-27 supplement minus vitamin A (Invitrogen Cat. #12587010), 1% P/S, 1% glutamine, and 1% sodium pyruvate. Prior to immediate use in culture, GNS media is supplemented with 20 ng/mL EGF (Peprotech Cat. #AF-100-15), bFGF (Peprotech Cat. #AF-100-18B), and PDGF-AA (Peprotech Cat. #AF-100-13A). Similarly with the GBM PDX cell lines, cell culture plates were incubated overnight at 4 °C with DPBS (with calcium and magnesium) and 10% laminin (Gibco^TM^ Cat. #23017015) prior to use. <u>Jurkat76 cell culture:</u> Jurkat76 cells were used as the TCR α- and β-negative human T-cell derivative that allowed for noncompeting introduction of exogenous TCRs.^7^ CD8+ Jurkat76 cells were cultured in RPMI supplemented with 10% fetal bovine serum and 1% P/S. <u>T2 cell culture:</u> T2 cells were used in the study to monitor immune cell response to the exogenous antigen of interest in a non-competitive environment. T2 cells are deficient in a peptide transporter involved in antigen processing (TAP), and as such, induction of these cells with exogenously administered peptides allows for their association and presentation by MHC class I molecules, HLA-A*0102 in particular.^8,9^ We cultured T2 cells in IMDM medium supplemented with 20% FBS. <u>COS7 and K562 cell culture:</u> We opted to use COS7 (ATCC Cat. #CRL-1651) and K562 (ATCC Cat. #CCL-243) cell lines as our respective primate and human artificial antigen presentation cell (aAPC) models.^10–12^ These cell lines do not express HLA molecules, which allows for the introduction of the HLA allele of interest. COS7 cells were cultured in DMEM medium supplemented with 10% FBS and 1% P/S. K562 cells were cultured in IMDM medium supplemented with 10% FBS and 1% P/S. <u>THP-1 cell culture:</u> THP-1 cells (ATCC Cat. #TIB-202) were used to investigate immune reactivity against neoantigen presentation by dendritic cells. THP-1 cells were cultured in RPMI-1640 supplemented with 10% FBS.

#### siRNA-mediated knockdowns of Splicing related genes

Cells were seeded in 2 mL of antibiotic free media in a 6-well plate at the following densities: GBM115 – 45,000 cells/well, SF10417 – 100,000 cells/well, and SF10602 – 100,000 cells/well. Twenty-four hours post seeding cells were transfected by adding 400 uL reaction containing serum free media, 2.0 uL DharmaFECT 1 reagent (Horizon, #T-2001-02), and their respective siRNA pools at a final concentration of 30 nM. Twenty-four hours post transfection, media was changed to complete media. At seventy-two hours post transfection, RNAs were isolated and purified via the Zymo Quick-RNA microprep kit (Zymo Research, #R1058).

### Reverse Transcription Quantitative PCR (RT-qPCR)

1000 ng of DNAse-treated RNA was converted to cDNA using the iScript cDNA synthesis kit (BioRad, #1708891). This cDNA was then diluted 1:3 using ultrapure, nuclease-free water, and 2 uL was used per qPCR reaction. Reverse transcription quantitative polymerase chain reaction (RT-qPCR) was performed using the Applied Biosystems POWER SYBR Green Master Mix (Applied Biosystems, #4367659). All samples were run in biological triplicates, with technical triplicates for each biological triplicate using the Quantstudio 5 (Thermo Scientific) and all gene expression data were normalized to the housekeeping gene *GUSB.* The cycling protocol is as follows: 2 minutes at 50 °C, 10 minutes at 95 °C, followed by 40 cycles of 95 °C for 15 s, and 60 °C for 60 s. Dissociation curves were performed to confirm specific product amplification. Primer sequences corresponding to each gene for the mRNA expression analysis were designed using NCBI Primer.

### Amplicon-sequencing for validation of neojunction expression

RNAs from respective cell lines were isolated and purified via the Zymo Quick-RNA microprep kit (Zymo Research, #R1058). 1000 ng of DNAse-treated RNA was converted to cDNA using the iScript cDNA synthesis kit (BioRad, #1708891). This cDNA was then diluted 1:3 using ultrapure, nuclease-free water, and 2 uL was used per PCR reaction. 16 reactions were carried out per amplicon per cell line using Q5 High-Fidelity 2x master mix (NEB, #M0492L) with primers containing partial Illumina adapters. Reaction mixtures were set up according to manufacturer guidelines. These products were then purified by separation on a 1.0% agarose gel at 100 volts constant for 1 hour and were then purified via the Monarch DNA gel extraction kit (NEB, #T1020L). Purified products were quantified with qubit high sensitivity dsDNA kit (Invitrogen, #Q32851) and prepared and submitted according to Azenta life sciences (Genewiz) guidelines for amplicon-sequencing.

### In-vitro sensitization of healthy donor PBMCs

HLA-A*02:01:01-positive PBMCs were purchased from StemExpress in either fresh or cryopreserved format. Fresh PBMCs (StemExpress Cat. #LE001F) of approximately 1x10^9^ cells were immediately proportioned into aliquots of 3 x 10^8^ cells and cryopreserved in liquid nitrogen, with one aliquot actively used for downstream IVS. Cryopreserved PBMCs (StemExpress Cat. #PBMNC300C) totaling approximately 3x10^8^ cells per cryovial were used in one vial per IVS procedure. PBMCs were thawed with 1:1000 Benzonase:RPMI (Sigma Aldrich Cat. #E8263). The CD14+ population was isolated from the PBMCs using CD14+. Miltenyi microbeads (Miltenyi Biotec Cat. #130-050-201) per manufacturer’s instructions. The CD14-flowthrough was cryopreserved for 6 days prior to naïve CD8+ T-cell isolation. Isolated CD14+ cells were cultured in CellGenix GMP DC medium (CellGenix Cat #20801-0500) supplemented with 1% human serum (Sigma Aldrich Cat #H6914), 1% P/S, 1000 U/mL recombinant human IL-4 (Peprotech Cat. #200-04), and GM-CSF (Peprotech Cat. #300-03) in non-treated 24-well plates at a seeding density of 5x10^5^ cells per well. On Day 3, recombinant human IL-4 and GM-CSF (1000 U/mL each) is added to the DC culture. On Day 5, DC culture is matured with 250 ng/mL LPS (Sigma Aldrich Cat. #L6529) in addition to supplementation of recombinant human IL-4 and GM-CSF (1000 U/mL each). Naïve CD8+ T-cells were isolated from the thawed CD14-population on Day 6 using the EasySep Human Naïve CD8+ T Cell Isolation Kit (STEMCELL Technologies Cat. #19258) as per manufacturer’s instructions. Isolated naïve CD8+ T-cells were cultured in X-Vivo 15 medium (Lonza Cat. #04-418Q) supplemented with 5% human serum, 1% P/S, and 10 ng/mL of recombinant human IL-7 (Peprotech Cat. #200-07) in 48-well plates at a seeding density of 5x10^5^ cells per well. On Day 8, adherent matured DCs were harvested from the plate using cold PBS. The collected DCs (1x10^6^ cells/mL) were exogenously pulsed with either 1 μM of the neoantigen peptide, influenza peptide, or no peptide for 1 hour at 37 °C. The peptide-pulsed or non-pulsed DCs were then co-cultured with naïve CD8+ T-cells at an optimal DC:T-cell ratio of 1:4 in 48-well plates. The co-culture was maintained with X-Vivo 15-medium supplemented with 10 ng/mL of recombinant human IL-7, 10 ng/mL recombinant human IL-15 (Peprotech Cat. #200-15), and 60 ng/mL of recombinant human IL-21 (Peprotech Cat. #200-21) for 10 days with IL-7 and IL-15 restimulation every 2 days. Cells were reseeded into subsequent 24-well, 12-well, and 6-well plates based on confluency. This concludes the first cycle of IVS of the neoantigens and influenza peptides. On Days 19 and 29, sensitized-CD8+ T-cells are reintroduced to a second and third round of stimulation with newly-pulsed DCs, and the co-culture is maintained for 10 additional days until the end of the second and third cycle of IVS. Immunogenic cytokine assays were performed at the end of the second and third cycles of IVS to determine whether a peptide-reactive T-cell population has expanded.

### Mutation-specific ELISA screen

Aliquots containing CD8^+^ T-cells from individual parent IVS wells were harvested and split equally into 96-well plate daughter wells containing 1x10^5^ cells per well. Daughter wells in triplicate were stimulated with T2 cells pulsed with either the neoantigen peptide of interest, control peptide, no peptide, or no T2 cells at all for 16 hours at an effector-to-target (E:T) ratio of 1:1. T2 cells were pulsed with 1 pM to 1 μM of the neoantigen peptide of interest, control peptide, or no peptides for 1 hour at 37 °C. Influenza-reactive T-cells were co-cultured against influenza peptide-pulsed T2 cells as a positive control. Co-culture supernatant was collected and diluted for use in IFNγ (BD Biosciences Cat. #555142) and TNFα (BD Biosciences Cat. #555212) ELISAs as per manufacturer’s instructions. ELISA readouts were performed on the Epoch Microplate Spectrophotometer (BioTek Instruments) using the BioTek Gen5 Data Analysis software (version 1.11). Wells with significantly increased expression levels of IFNγ and TNFα were selected for downstream single-cell immune profiling using single-cell RNA and V(D)J sequencing.

### Single-cell immune profiling

Once an expanded neoantigen-reactive CD8^+^ T-cell population from IVS is identified, single-cell RNA and V(D)J sequencing were performed using the 10x Genomics platform. Prior to sequencing, CD8^+^ T-cells from the expanded neoantigen-reactive (ELISA screen-positive) wells were harvested and co-cultured with T2 cells pulsed with 1 μM of the neoantigen peptide of interest, a control peptide, or no peptides at an E:T ratio of 1:1. One co-culture replicate was performed for 3 hours for single-cell RNA sequencing analysis, and another was performed for 16 hours for IFNγ and TNFα ELISA confirmation. The final cell concentration was adjusted to approximately 1x10^4^ cells/μL with an initial cell viability of at least 90% to maximize the likelihood of achieving the desired cell recovery target. Independent CD8^+^ T-cell and non-pulsed T2 single cultures were sequenced alongside the co-culture conditions for differentiating cell types in the downstream single-cell sequencing analysis. The Chromium Next GEM Single Cell 5’ Reagent Kit v2 (Dual Index) (10xGenomics, Cat. #CG000331) was used for preparation for single-cell sequencing analysis. Gel bead in emulsions (GEMs) were generated by combining the single cell 5’ gel beads, partitioning oil, and the master mix containing the cells onto the Chromium Next GEM Chip K. Cell lysis and barcoded reverse transcription of RNAs in all single cells were finished inside their corresponding GEM. Barcoded cDNA product was recovered through post-GEM-RT cleanup and PCR amplification. cDNA quality control and quantification were performed on the Fragment Analyzer System (Agilent Technologies). 50 ng of cDNA was used for the construction of the 5’ gene expression library, and each sample was indexed by a Chromium i7 Sample Index Kit. This process was performed on an Illumina NovaSeq 6000 sequencer at the UCSF Institute of Human Genetics (IHG) with a minimum of 20,000 read pairs per cell for the 5’ Gene Expression library. The enriched product was measured by the Fragment Analyzer System. 50 ng of enrichment TCR product was used for library construction. Single-cell V(D)J enriched libraries were subsequently sequenced on the Illumina NovaSeq 6000 with a minimum of 5,000 read pairs per cell for the V(D)J library. Cell Ranger 7.0.0 (10x Genomics Cloud Analysis) was used to pre-process raw single-cell RNA sequencing and identifying V(D)J clonotypes. The annotation files ‘vdj_GRCh38_alts_ensembl-3.1.0-3.1.0’ and ‘GRCh38-3.0.0’ were used for demultiplexing cellular barcodes, performing read alignments, and generating feature-barcode matrices. Only cells for which clonotype information was available were retained for downstream analysis. Single-cell gene expression and corresponding V(D)J sequences of candidate T-cell clonotypes were analyzed on the Loupe V(D)J browser. Single cells with detectable *CD8A* expression were specifically isolated and characterized as the CD8^+^ T-cell population and subsequently grouped according to their TCR clonotypes. To identify T-cell clonotypes associated with a neoantigen-specific response, we selected expanded TCR clonotypes with significantly increased levels of *IFNG*, *TNF*, and *GZMB* expressions in the T-cell:neoantigen-pulsed T2 condition compared to the T-cell:control-pulsed T2 and T-cell:non-pulsed T2 conditions.

### HLA typing

OptiType 1.3.1 was used for genotyping HLA alleles from available WES data available for glioma cell lines with default parameters.

### Plasmids and peptides

HLA-A*02:01 and neojunction-derived gene sequences were all synthesized and cloned into the pTwist Lenti SFFV Puro WPRE vector (Twist Biosciences). Constructs encoding full-length and truncated multi-mer versions of the wildtype and mutant *GNAS* and *RPL22* sequences were generated. TCR α/β was synthesized and cloned into the pTwist Lenti SFFV vector (Twist Biosciences). HPLC grade neojunction-derived neoantigen peptide multi-mers (>95%) were manufactured by TC Laboratories.

### Lentiviral transduction

HEK293T cells were plated in 6-well culture plates at a density of 1x10^6^ cells per well with 2 mL DMEM medium supplemented with 10% FBS without antibiotics. After approximately 18 to 24 hours or at 90% confluency, HEK293T cells were transfected with the expression construct, see above, and lentiviral packaging plasmids, pMD2.G (Addgene, #12259) and psPAX2 (Addgene, #12260). <u>TCR α/β transduction:</u> 1.0 μg TCR α/β transfer plasmid, 0.75 μg psPAX2, and 0.25 μg pMD2.G were combined with 200 μL Opti-MEM (Thermo Fischer Scientific Cat. #31985062). 6 μL of Xtremegene HP was added to this mixture and complex formation was allowed to occur for 15 minutes at room temperature at which point this reaction mixture was added to corresponding HEK293T cells. Transfection medium was replaced with fresh DMEM media after 24 hours. Viral supernatant was collected after 48 hours, and a functional virus titer was performed on 6-well plates seeded with Jurkat76/CD8 cells or PBMC-derived CD8^+^ T-cells at 60-70% confluency. Viral transduction was performed with 3-fold serial dilutions of the virus stock supplemented with polybrene at a final concentration of 4 μg/mL. Media was changed 24 hours following viral transduction. Cells were assessed for transduction efficiency after 3-4 days by measuring surface expression of TCR α/β and CD3 by fluorescence-activated cell sorting (FACS) analysis. Cells demonstrating high double-positive expression of TCR α/β and CD3 were flow-sorted and maintained for downstream co-culture and immunogenicity assays. <u>HLA and neoantigen transduction:</u> Constructs expressing HLA-A*02:01 were linearized and restricted with BamHI and XhoI (New England Biolabs) and purified using the Zymoclean Gel DNA Recovery Kit (Zymo Research Cat. #D4007). The HLA-A*0201 sequence was then ligated into a lentiviral construct downstream of an EF1A-core promoter and upstream of an IRES followed by a Blasticidin resistance gene. 1.0 μg of either HLA-A*02:01 or neoantigen transfer plasmid, 0.75 μg psPAX2, and 0.25 μg pMD2.G were combined with 200 μL Opti-MEM (Thermo Fischer Scientific Cat. #31985062). 6 μL of Xtremegene HP was added to this mixture, and complex formation was allowed to occur for 15 minutes at room temperature at which point this reaction mixture was added to corresponding HEK293T cells. As stated above, neoantigen constructs encode either the full-length or truncated version of the neojunction-derived peptide. The transfection medium was replaced with fresh DMEM media after 24 hours. HLA-A*02:01 lentiviral transduction and screening was performed first prior to neoantigen lentiviral transduction and screening for streamlined drug selection. Viral supernatant was collected after a subsequent 48 hours, and a functional virus titer was performed on 6-well plates seeded with COS7 or K562 cells at 60-70% confluency. Viral transduction was performed with 3-fold serial dilutions of the virus stock supplemented with 4 μg/mL polybrene. Media was changed 24 hours following viral transduction and replaced with complete media supplemented with blasticidin. Cells were assessed for transduction efficiency after 3-4 days by drug screening. HLA-A*02:01-transduced APCs were cultured in medium treated with 10 μg/mL Blasticidin for approximately 7 days before assessing for cell viability across titers. Neoantigen-lentiviral transduction is subsequently performed, and APCs transduced with both HLA-A*02:01 and neoantigen-expressing constructs are then cultured in medium treated with 3 μg/mL puromycin for approximately 7 days. Cell viability was assessed afterwards across all titer conditions. Cells were assessed for transduction efficiency after 3-4 days by measuring surface expression of HLA-A2 fluorescence-activated cell sorting (FACS) analysis.

### Dose-dependent assessment of TCR reactivity against neoantigen

Specificity of neoantigen-reactive CD8^+^ T-cells and TCR-transduced T-cells was assessed by human IFNγ (BD Biosciences Cat. #555142), IL-2 (BD Biosciences Cat. #555190), and TNFα ELISA (BD Biosciences Cat. #555212). Assessment of TCR recognition against exogenously introduced neoantigen peptides presented by MHC-I molecules was conducted by co-culturing T-cells with peptide-pulsed T2 cell conditions. T2 cells are either pulsed with neoantigen peptide of interest at a concentration between 1 pM through 1 μM, decoy peptide, or no peptides for 1 hour at 37°C. Influenza-reactive T-cells are co-cultured against influenza peptide-pulsed T2 cells as a positive control. T-cells and T2 cells were co-cultured in a 96-well round-bottom plate at a concentration of 1 x 10^5^ of each cell type in 200 μL of medium for 16 hours. Supernatant was collected and diluted for cytokine release assays per manufacturer’s instructions. ELISA assay readouts were performed on Epoch Microplate Spectrophotometer (input wavelength 450 nm and output wavelength 570 nm) using the BioTek Gen5 Data Analysis software. To characterize the dose-dependent activation of the TCRs in transduced triple-reporter Jurkat76/CD8 cells, we performed flow analysis to assess the level of expression of NFAT-GFP, NFκB-CFP, AP-1-mCherry following 16 hours of co-culture. Similarly, reactivity of TCR-transduced PBMC-derived CD8^+^ T-cells was evaluated by flow analysis following anti-CD107a (BioLegend, Cat #328620) and anti-CD137 antibody (4-1BB; Biolegend Cat #309804) staining.

### In vitro transcription (IVT) synthesis of mRNA

All constructs were subcloned into pcDNA3.1 (Invitrogen, 2520855) and linearized by XhoI restriction enzyme with the plasmid DNA template transcribed downstream from the bacteriophage T7 promoter sequence. For long (> 0.5 kb) and short (< 0.5 kB) transcripts, 1 μg and 0.5 μg of template were used, respectively. Reactions were assembled at room temperature using the mMESSAGE mMACHINE T7 Transcription Kit as per manufacturer’s instructions (Invitrogen, 2582905) and incubated at 37°C for 1 hour for long transcripts and 16 hours for short transcripts. Following DNase treatment, a Poly(A) tailing reaction was performed for 1 hour according to the HiScribe T7 ARCA manual (NEB, E2060S). Subsequently, the synthesized mRNA was purified by LiCl precipitation using 70% DEPC-based ethanol. Synthesized mRNA was heat-shocked (70[, 5mins) with the formaldehyde loading dye to verify quality via gel electrophoresis.

### mRNA transfection of HLA-A*02:01, truncated neoantigen, and full-length neojunction-encoding mRNA

Transfection of IVT-synthesized mRNA into COS7 and K562 cells was performed with electroporation using the Neon Transfection System 100 μL Kit (Invitrogen, MPK10096) per manufacturer’s instructions. 1 x 10^6^ COS7 and K562 cells were washed and resuspended with 100 μL of Neon^TM^ Resuspension Buffer. 5 μg of HLA-A2 and 5 μg of candidate (either the truncated neoantigen sequence or the full-length neojunction sequence) mRNA were added into the cell solution. Electroporation was performed on the Neon NxT Electroporation System (Invitrogen, NEON1). Electroporation of COS7 cells was performed with the following optimized conditions: pulse voltage of 1200 V, width of 30 ms, and 2 pulses. Electroporation of K562 was performed with the following optimized conditions: pulse voltage of 1450 V, width of 10 ms, and 3 pulses. Transfected cells were immediately transferred into warm RPMI with no antibiotics. Aliquots of transfected cells were retained for validation of HLA-A2 expression by staining with HLA-A2 monoclonal antibody (BB7.2, Thermo Scientific, 17-9876-42) and subsequent flow cytometry analysis.

### Evaluation of TCR specificity against endogenously processed and MHC-I presented neoantigen

Characterization of neoantigens that are endogenously processed and presented by surface MHC-I is conducted by co-culturing HLA-A*02:01/neoantigen-transfected COS7 or K562 cells with TCR-transduced T-cells. Similarly, T-cells and COS7/K562 cells were co-cultured in a 96-well flat-bottom plate at a concentration of 1 x 10^5^ of each cell type in 200 μL of medium for 16 hours. Supernatant was collected and diluted for cytokine release assays per manufacturer’s instructions, and cytokine release levels were assessed with the Epoch Microplate Spectrophotometer and BioTek Gen5 Data Analysis software. In all cytokine release assay experiments, maximum cellular cytokine release per well was determined by the addition of 0.2 μL Cell Activation Cocktail (without Brefeldin A) (BioLegend Cat. #423302) per 100 μL cell solution. Evaluation of endogenously processed and presented neoantigens in glioma cell lines was performed by co-culturing TCR-transduced triple-reporter Jurkat76 cells with glioma cells at a 1:1 E:T ratio (1 x 10^5^ per well in a 96-well plate). Flow analysis was performed to assess the level of expression of NFAT-GFP, NFκB-CFP, AP-1-mCherry following 16 hours of co-culture.

### HLA-IP and LC–MS/MS

COS-7 cells were co-electroporated with 10 μg of each mRNA encoding HLA-A*02:01 allele and the full-length coding sequence of the mutated GNAS or RPL22 using the Neon Transfection system (100-μl tip, setting: 1,050 V/10 ms/2 pulses). 20[×[10^6^ cells were electroporated per condition and plated in six-well non-TC plates overnight. For the GMB115 cell line sample, approximately 100×10^6^ cells were used. Cells were harvested by incubating with 1 mM EDTA (Millipore Sigma) for 10[minutes at 37 °C. For the immunoprecipitation experiments, cells were lysed in 8[ml of 1% CHAPS (Millipore Sigma) for 1 hour at 4°C, lysates were then spun down for 1 hour at 20,000g and 4°C, and supernatant collected. For the affinity-column based immunopurification of HLA-I ligands, 40 mg of cyanogen bromide– activated–Sepharose 4B (MilliporeSigma) was activated with 1 mM hydrochloric acid (MilliporeSigma) for 30 minutes. Subsequently, 1 mg of W6/32 antibody (Bio X Cell) were coupled to Sepharose in the presence of binding buffer (150 mM sodium chloride, 50 mM sodium bicarbonate, pH 8.3; sodium chloride) for 2 hours at room temperature. Sepharose was blocked for 1 hour with glycine and washed 3 times with PBS. Supernatants of cell lysates were run over the affinity of column through peristaltic pumps at 6 mL/min flow rate overnight at 4°C. HLA complexes and binding peptides were eluted from the column five times using 1% TFA. Peptides and HLA-I complexes were separated using C18 columns (Sep-Pak C18 1[cc Vac Cartridge, 50 mg of sorbent per cartridge, 37–55-μm particle size, Waters). C18 columns were pre-conditioned with 80% ACN (Millipore Sigma) in 0.1% TFA and equilibrated with two washes of 0.1% TFA. Samples were loaded, washed twice with 0.1% TFA and eluted in 300[μl of 30%, 40% and 50% acetonitrile in 0.1% TFA. All three fractions were pooled, dried down using vacuum centrifugation and stored at −80 °C until further processing. HLA-I ligands were isolated by solid-phase extractions using in-house C18 mini-columns. Samples were analyzed by high-resolution/high-accuracy LC–MS/MS (Lumos Fusion, Thermo Fisher Scientific). COS-7 samples we run at DDA mode while GMB115 samples at DIA. MS and MS/MS were operated at resolutions of 60,000 and 30,000, respectively. Only charge states 1, 2 and 3 were allowed. The isolation window was chosen as 1.6 Thomson, and collision energy was set at 30%. For MS/MS, maximum injection time was 100 ms with an automatic gain control of 50,000. MS data were processed using FragPipe. Protein FDR was set at 1%. Oxidization of methionine, phosphorylation of serine, threonine and tyrosine, as well as N-terminal acetylation were set as variable modifications for all samples. Samples were searched against a database comprising UniProt Cercopithecus aethiops or Uniprot Human reviewed proteins supplemented with human HLA-A*02:01 allele sequence, mutRPL22 and mutGNAS, as well as common contaminants.

### Characterization of CD8+ T-cell-mediated anti-tumor reactivity

To determine whether TCR-transduced T-cells were capable of mounting an anti-tumor response, TCR-transduced Jurkat76/CD8 or PBMC-derived CD8^+^ T-cells were co-cultured with patient-derived GBM or LGG cell lines. CD8+ T-cells were isolated from healthy donor-derived PBMCs using the EasySep™ Human CD8^+^ T Cell Isolation Kit (STEMCELL Technologies, Cat. # 17953). CD8^+^ T-cells were then activated with Dynabeads™ Human T-Activator CD3/CD28 for T Cell Expansion and Activation (Thermo Scientific, Cat. #11161D) at a concentration of 25 μL/1 x 10^6^ cells. CD8^+^ T-cells were cultured for 7 days with IL-7 (30 μL/1 x 10^6^ cells) supplemented every 2 days. CD8^+^ T-cells were then LV transduced with neoantigen-specific TCRs with a hybridized murine TCR constant region using the above transduction procedure. This additional step removes the likelihood of TCR α-chain and β-chain mispairing and allows us to evaluate TCR-transduction efficiency by staining with anti-murine TCR constant region antibody (Clone H57-597; BioLegend Cat. #109208). Flow sorting was performed to isolate highly-transduced CD8^+^ T-cells by selecting for cells stained strongly with anti-CD3 and anti-murine TCR constant region antibody. Sorted transduced CD8^+^ T-cells were expanded for 7 days before use in co-culture assays. Killing assays were performed using an xCELLigence RTCA S16 Real-Time Cell Analyzer. Tumor cells were cultured in media pre-treated with 100 ng/mL IFNγ (Peprotech, Cat. #300-02) for 48 hours and washed twice with PBS prior to seeding. 1 x 10^4^ tumor cells were plated per well in a 96-well E-plate (Agilent), and impedance is read for 16 hours during incubation. TCR-transduced CD8^+^ T-cells were introduced to each well at E:T ratio of either 1:1 or 2:1, and tumor-specific killing is measured by changes in cell index over 24-48 hours.

### Identification of MHC-I restricted CD8^+^ T-cell mediated reactivity against neoantigens

Evaluation of MHC-I restricted reactivity was performed by perturbing TCR and HLA:peptide interaction with the introduction of anti-MHC-I antibodies. In dose-dependent immunogenicity assays, T2 cells at a concentration of 1 x 10^5^ tumor cells per well of a 96-well plate were washed twice with PBS and incubated for 30 min with blocking anti-MHC-I antibody (50 μg/well; clone W6/32, Bio X Cell, Cat. #BE0079) or isotype control (50 μg/well; Bio X Cell, Cat. #BE0085) at a total volume of 100 μL. Without any additional washes, T-cells were added in to achieve a final volume of 200 μL. In tumor-killing assays, tumor cells were added to each well of a 96-well E-plate in a total volume of 50 μL for initial seeding. Anti-MHC-I antibody or isotype control (50 μg/well) is added to each well 30 minutes prior to the addition of T-cells to reach a total volume of 100 μL. T-cells were added in each well to achieve a final volume of 200 μL, and impedance was measured for the following 24-48 hours.

### FACS analysis and antibodies

TCR-transduced cell lines were stained with anti-human TCR α/β (Clone IP26, BioLegend Cat. #306717) and anti-human CD3 antibody (Clone HIT3a, BioLegend Cat. #300307) to assess the surface-level expression of the transduced TCR. CD8^+^ T-cells were stained with anti-CD107a (BioLegend, Cat #328620) and anti-CD137 antibody (4-1BB; Biolegend Cat #309804) to assess CD8^+^ T-cell degranulation and TCR activation, respectively. Viability of cells were assessed with the Zombie Green^TM^ Fixable Viability Kit (BioLegend, Cat. #423111) APCs and patient-derived glioma cell lines were stained with HLA-A2 monoclonal antibody (Clone BB7.2, Thermo Fisher Scientific Cat. #17-9876-42). Approximately 1x10^6^ cells per 100 μL FACS buffer (PBS supplemented with 1% BSA (Sigma Aldrich Cat. #L6529)) is incubated with one test volume of antibody for 20 minutes as indicated by the manufacturer. Stained cells were washed once with FACS buffer before resuspension to a concentration of 4x10^5^ cells per 100 μL FACS buffer. Cells were then analyzed with the Attune NxT flow cytometer (Thermo Fischer Scientific).

### Gene set enrichment analysis

Differential gene expression of TCGA, GTEx, and UCSF GBM/LGG RNA-sequencing was performed and quantified using DESeq2.^13^ Only genes with an absolute fold change > 1.5 and a Benjamini-Hochberg adjusted *P* value <0.05 called by DESeq2 were considered to be differentially expressed.^14^ Pre-ranked gene set enrichment analysis (GSEA)^15^ was carried out by ranking genes with the product of their fold-change sign and the -log10(adjusted *P* value). <u>Disease subtype-</u> <u>specific differential gene analysis:</u> GSEA comparison was performed between *IDH1* mutation subtypes (wildtype *IDH1* and mutant *IDH1*) as well as glioma disease subtypes (wildtype *IDH1* glioblastoma, mutant *IDH1* astrocytoma, and mutant *IDH1* oligodendroglioma). Splicing-related gene sets were selected based on keyword search, and gene sets with an adjusted *P* value <0.05 when comparing two groups are considered differentially enriched. Unbiased hierarchical clustering of differentially enriched gene sets allows the characterization of subgroup-specific upregulated genes. <u>Neojunction load-specific differential gene analysis:</u> TCGA LGG and GBM samples were ranked according to the total putative neojunctions expressed per sample. High (NJ_HI_) and low neojunction load (NJ_LO_) samples within each disease subtype were characterized as the upper and lower 0.10 percentile of ranked samples, respectively. GSEA is carried out between the NJ_HI_ and NJ_LO_ samples of each disease subgroup. Gene sets with a unidirectional fold-change and adjusted *P* value <0.05 were considered to be enriched gene sets associated with neojunction load. Splicing-related gene sets were selected based on keyword searches. Leading edge genes shared across all disease subgroups within the same gene set are defined as enriched genes associated with neojunction load.

### Neojunction and splicing-related gene correlation analysis

Selection of mutant *IDH1* upregulated genes was determined by splicing-related genes expressed with a significant (*p* < 0.05) log2fold increase of 1.5 in mutant *IDH1* cases when compared to their wild-type counterpart. Selection of splicing-genes affected by oligodendroglioma-specific loss of chromosomes 1p/19q was determined by chromosome 1p/19q splicing-related genes expressed with a significant (*p* < 0.05) log_2_fold decrease of 1.5 in IDH-O cases compared to both IDH-A and IDH-wt cases. Splicing-related genes that were selected for *in vitro* validation was chosen based on previously reported confirmation of aberrant splicing due to their dysregulated expression.^16–20^ To determine correlation factors between each of the identified public neojunctions with each splicing-gene of interest, we performed a Pearson correlation analysis against each neojunction and splicing-related gene pair. Neojunctions with the highest positive correlation score against the select mutant *IDH1* upregulated genes (*CELF2*, *ELAVL4*) averaged across all three glioma subtypes were tested in downstream qPCR assays. Similarly, neojunctions with the most negative correlation score against select chromosome 1p or 19q splicing-related genes downregulated in *IDH1*mut-O cases (*SNRPD2*, *SF3A3*) averaged across all three glioma subtypes were also tested in downstream qPCR assays.

## QUANTIFICATION AND STATISTICAL ANALYSIS

All statistical analysis was performed in base R.

